# Saturation of influenza virus neutralization and antibody consumption can both lead to bistable growth kinetics

**DOI:** 10.1101/2020.03.02.973958

**Authors:** Shilian Xu, Ada W. C. Yan, Heidi Peck, Leah Gillespie, Ian G. Barr, Xiaoyun Yang, Stephen J. Turner, Celeste M Donato, Tonghua Zhang, Moshe Olshansky, Dhanasekaran Vijaykrishna

## Abstract

Influenza virus is a major human health threat. Neutralizing antibodies elicited through prior infection or vaccination play an irreplaceable role in protection from subsequent infection. The efficacy of antibody-dependent vaccines relies on both virus replication and neutralization, but their quantitative relationship was unknown. Here we use mathematical models to quantitatively investigate viral survivability determined by antibody concentration and inocula size. We performed focus reduction assays for 49 seasonal influenza A/H3N2 viruses circulating during 2017–2019 against influenza antisera raised in ferrets, and find that the antibody consumption rates of individual reactions were either small or large, and this was strongly positively correlated with virus saturation. Regardless of antibody consumption rate, virus-antibody interactions always lead to antibody-induced bistable viral kinetics. As a result, at a specific interval of antibody concentration, small viral inocula are eliminated but not large virus inocula, which is triggered by saturated virus neutralization or antibody consumption. Our finding highlights virus-antibody interaction with different antigenic properties, thereby explaining commonly observed influenza re-infection and enhancing vaccine efficiency.

## Introduction

Influenza virus is a major human health concern responsible for an estimated 290,000–650,000 deaths annually, particularly in high-risk groups such as pregnant women, immunocompromised patients and individuals with comorbidities [1–4]. Antibody-mediated immunity provides a robust and relatively long-lived protection from severe disease against virus strains [5–7]. However, exposure to influenza through infection or vaccination does not guarantee exemption from sequential infection due to the same influenza antigenic variant. Influenza viruses are often isolated from vaccinated individuals and re-infection due to the same influenza antigenic variant is commonly observed in the laboratory [8], and during seasonal influenza epidemics (for example 17% re-infection rate of A/H3N2 in 1970, 23% of A/H3N2 in 1976, 20% of A/H1N1 in 1980, 32% of A/H3N2 in1983 and 25% of influenza B [9–13]). Trivalent or quadrivalent influenza vaccine can significantly reduce disease severity, but can only protect 20%-60% individuals who receive them [14, 15]. These results show that pre-existing antibody mediated immunity cannot ensure exemption of future infection [16], hampering progress towards developing a efficacious influenza vaccine.

Serological assays aim to quantify the prevalence of neutralising antibodies against specific influenza strains, primarily through the hemagglutinin inhibition (HI) assay. Serum HI antibody titres of 40 units (i.e. 1:40 or lesser dilutions) against an influenza antigenic variant is assumed to reduce risk of infection by 50% in the population; and is accepted as a reasonably accurate correlate of protection by many regulatory agencies [17, 18]. While the HI assay is important to determine the levels of antibodies to influenza in a sample and quantify antigenic evolution, they do not inform the antibody-virus kinetics as the amount of neutralized virus is not quantified, leaving gaps in our understanding of virus neutralization kinetics. Further, the HI assay can also be affected by variation in serum potency and virus binding activity [19].

Simple compartmental models, particularly the target cell-infected cell-virus (TIV) model, have been used to provide a great deal of understanding of the within-host kinetics of influenza virus infection [20–25], and have played a key role in quantifying pre-existing immunity [21, 25–27]. However, in existing models with virus-antibody interaction, the rate of virus neutralization is assumed to be directly proportional to the product of antibody concentration and viral titre, allowing infinite increases in neutralisation rate with increase of viral titre or antibody concentration [20, 21, 23, 24, 26, 27]. Since, both *in vivo* and *in vitro*, the rate of antibody binding saturates at a concentration higher than that required for neutralization, for a fixed viral titre, the rate of virus neutralisation increases and is saturated with increases of antibody concentration (Fig. 1b). Saturation commonly occurs in biological systems; whose effects are innately nonlinear in proportion to reactant concentrations[28–30]. Crucially, saturation always leads to a bistable behaviour – biologically known as inoculum effect – where two possible outcomes are possible [31] (Fig. 1c). It is not known whether saturation has a bistable effect on the survival or eradication of influenza following infection in the presence of antibodies.

**Fig. 1.**
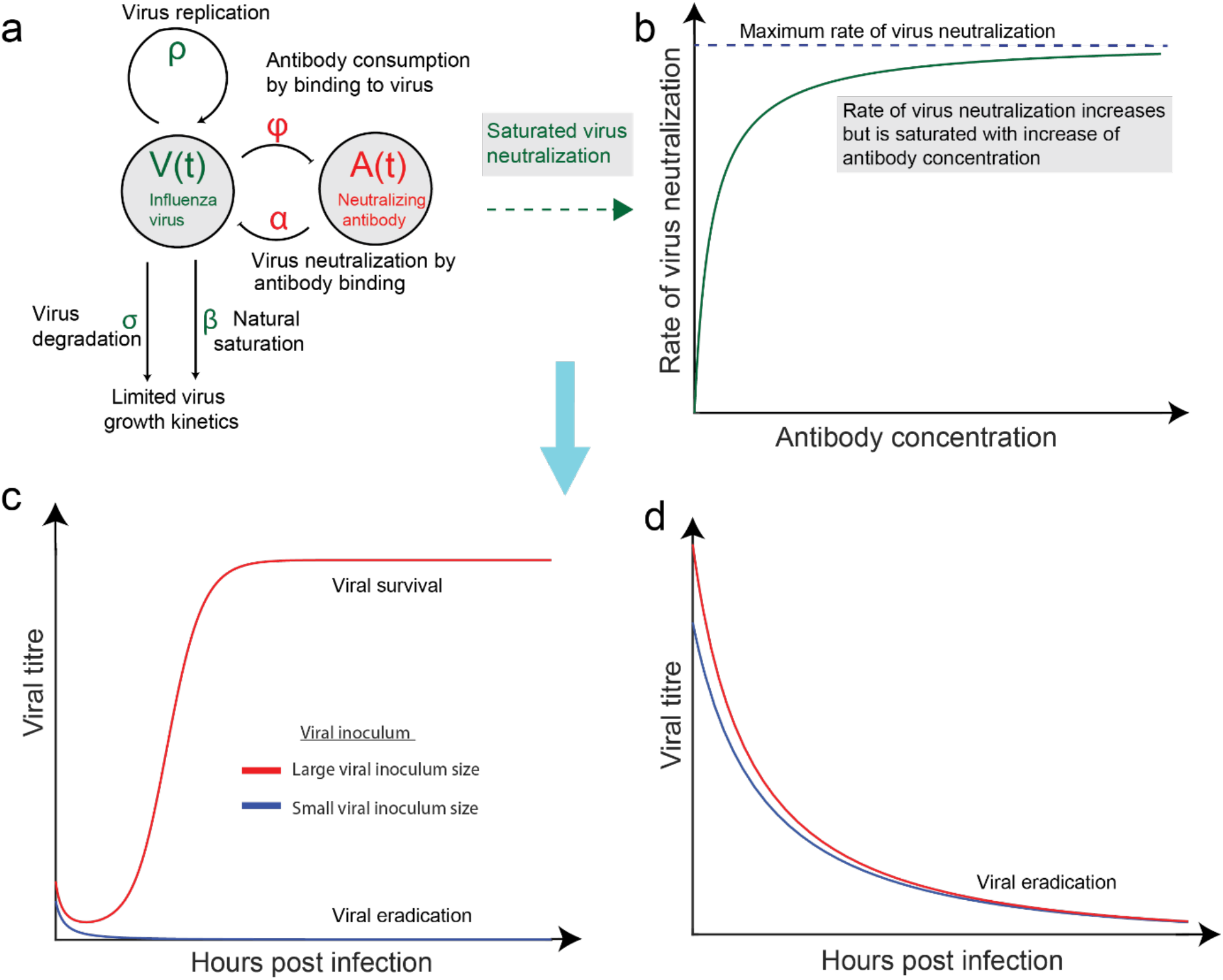
Schematic of viral kinetics with saturated virus neutralization to investigate qualitatively viral inoculum size on viral survival and eradication. A) Virus kinetic with saturated virus neutralization is divided into limited viral growth kinetics (virus replication, virus degradation and natural saturation on limited cell number) and virus neutralization by antibody binding (saturated virus neutralization and antibody consumption). B) Saturated virus neutralization refers that rate of virus neutralization increases but converges to maximum rate of virus neutralization with increase of antibody concentration. C) Neutralizing antibody induces bistable viral kinetics; large virus inocula survive and small virus inocula are inhibited. In this case, variability of virus neutralization can be innately biological features of the interaction between virus and antibody, rather than antigenic change driven by mutations. D) Neutralizing antibody induces monostable viral kinetics; influenza virus is inhibited independent of viral inoculum size. Then, variability of virus neutralisation is only caused by antigenic changes.

To understand how saturation of virus neutralization affects viral kinetics, we developed a class of two dimensional models that involve virus neutralisation estimated by performing focus reduction assays (FRA) of seasonal influenza A H3N2 viruses and virus growth kinetics. We found that the rate of virus neutralization saturated with increases of both viral titre or antibodies, and our models allowing saturated virus neutralization provided a robust and better fit than unsaturated neutralisation. Integrating influenza replication and neutralization parameters into the proposed deterministic models, we identified saturation of virus neutralization or antibody consumption lead to bistable viral growth kinetics in the presence of neutralizing antibodies. We also found a strong positive correlation between the antibody consumption and virus saturation, and that they could be categorised into two groups based on the rate of antibody consumption, information that could be utilised for vaccine preparation. Overall, our analysis reveal that antibody-induced bistable viral kinetics exist through saturated virus neutralization and antibody consumption. This shows even for virus-antibody pair that are well-matched, variability of virus neutralization can always occur, thereby explaining the occurrence of reinfections due to antigenically similar influenza strains.

## Results

### Virus neutralization is saturated with increase of both viral titre and antibody concentratio

Virus neutralization kinetics were quantified for 49 seasonal influenza A/H3N2 viruses circulating during 2014–2019 (HA clade 3C.2A [32]) against antisera raised in ferrets against eight reference viruses using the focus reduction assay (FRA), a neutralization assay based on immunostaining, allowing the estimating of total viral titre and neutralized viral titre before and during virus-antibody incubation (see *Methods*). For all viruses, the neutralized viral titre increased and converged to a total viral titre with increase of antibody concentration, independent of combination of influenza virus and antisera (Fig. 2a).

**Fig. 2.**
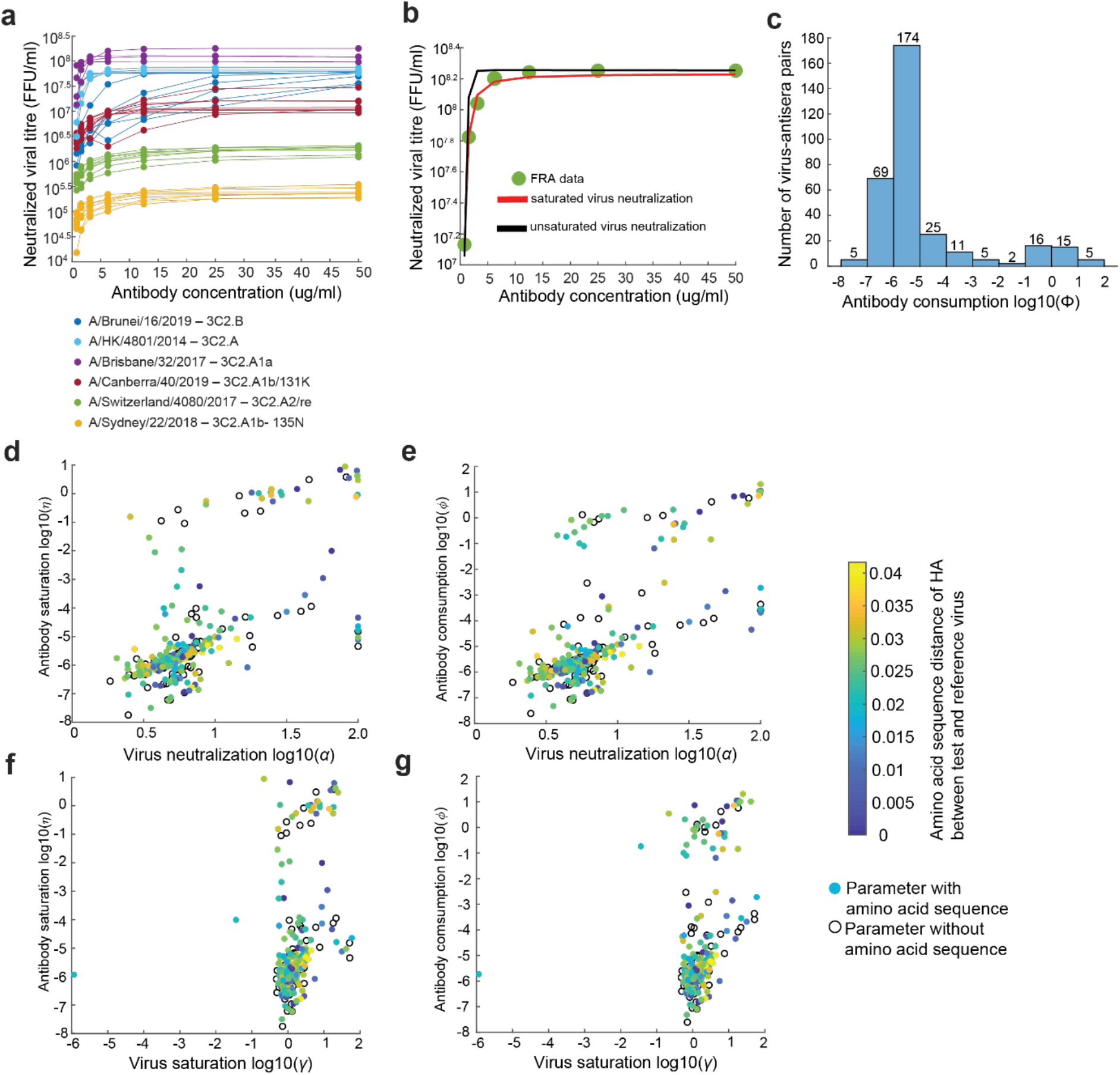
Analysis of FRA data arisen from 49 test virus against eight antisera. A) Neutralized viral titre increases and converges with increase of antibody concentration independent of A/Hong Kong/4801/2014 (3C2.A), A/Brunei/16/2019 (3C2.B), A/Switzland/4080/2017 (3C2.A2/re), A/Canberra/107/2019 (3C2.A1b/131K), A/Sydney/22/2018 (3C2.A1b/135N) and A/Brisbane/32/2017 (3C2.A1a). b) Saturated virus neutralization (red curve) fits better to FRA data (green cycle) than unsaturated virus neutralization (black curve). C) Histogram plotting antibody consumption against number of FRA dataset exhibits bimodal behaviours, one peak (majority of FRA data) locating on antibody consumption from 10^−7^ to 10^−5^ and another peak (minority of FRA data) locating on antibody consumption from 10^−1^ to 10^1^. This indicates FRA datasets are classified into two categories, large and small antibody consumption. D-g) Antibody saturation and antibody consumption are strongly positively correlated. D) Virus neutralization *log*_10_(*α*) is plotted against antibody saturation *log*_10_(*η*); e) virus neutralization *log*_10_(*α*) is plotted against antibody consumption *log*_10_(*φ*); f) virus saturation *log*_10_(*γ*) is plotted against antibody saturation *log*_10_(*η*); g) virus saturation *log*_10_(*γ*) is plotted against antibody consumption *log*_10_(*φ*).

To establish a quantitative relationship between neutralised virus titre and antibody concentration we developed two models of virus neutralisation, saturated (System 1, methods) and unsaturated (System 2, methods) to describe the rate of change of viral titres and antibody concentration (Illustrated in Fig. 1a). As expected the model allowing saturated virus neutralization better fit our FRA estimates than a model with unsaturated neutralisation commonly assumed in prior studies [20, 21, 24, 27] (Fig. 2b). To test for overfitting, we generated simulated data with noise following Gaussian distribution *N*(0,1) of different magnitudes and then fit saturated and unsaturated virus neutralization models (Section 2.2, *Supplementary material*), to find that the saturated neutralization model is more robust to tolerate noise than unsaturated virus neutralization despite containing two fewer parameters. As an example, virus neutralization parameters estimated for A/Canberra/40/2019 is shown in Fig 2c and Table 2, showing that the rate of virus neutralization increases but is saturated with increase of both antibody concentration and viral titer.

**Table 1.** Raw Focus Reduction Assay data generated using A/Canberra/40/2019 (FFU scale)

| Serum Dilution | Antisera arisen from reference virus |  |  |  |  |  |  |  |
| --- | --- | --- | --- | --- | --- | --- | --- | --- |
|  | A/Hong Kong/4801/2014 (Egg grown) | A/Newcastle/82/2018 (cell grown) | A/Newcastle/82/2018 (egg grown) | A/Sydney/22/2018 (cell grown) | A/Victoria/653/2017 (cell grown) | A/Victoria/653/2017 (egg grown) | A/Switzerland/8060/2017 (cell grown) | A/Switzerland/8060/2017 (egg grown) |
| 80 | 195 | 52 | 102 | 18 | 8 | 142 | 17 | 16 |
| 160 | 449 | 62 | 43 | 47 | 15 | 532 | 25 | 10 |
| 320 | 2278 | 493 | 339 | 144 | 37 | 1226 | 85 | 29 |
| 640 | 3405 | 1296 | 1019 | 576 | 131 | 1573 | 306 | 116 |
| 1280 | 3780 | 1791 | 1861 | 1456 | 578 | 1441 | 798 | 489 |
| 2560 | 4069 | 2136 | 1937 | 1424 | 1318 | 1173 | 1148 | 965 |
| 5120 | 4416 | 2226 | 2034 | 1545 | 1459 | 1481 | 1098 | 958 |
| 10240 | 4924 | 2606 | 2590 | 1905 | 1747 | 1890 | 1681 | 1492 |

**Table 2.** Estimated virus neutralization parameters obtained from Focus Reduction Assay using A/Canberra/40/2019

| Reference virus (cell/egg grown) | $\alpha$<br>(ml/FFU/hour<br>) | $\eta$<br>(ml/ug) | $\gamma$<br>(ml/FFU) | $\varphi$<br>(ml/ug/hour) | Category |
| --- | --- | --- | --- | --- | --- |
| A/Hong Kong/ 4801/201(egg) | 5.3848 | $1.91 \times 10^{-6}$ | 1.6052 | $2.49 \times 10^{-6}$ | Small |
| A/Newcastle/82/2018 (cell) | 6.5889 | $1.62 \times 10^{-6}$ | 1.4073 | $2.19 \times 10^{-6}$ | Small |
| A/Newcastle/82/2018 (egg) | 5.9281 | $1.28 \times 10^{-7}$ | 1.695 | $1.70 \times 10^{-7}$ | Small |
| A/Sydney/22/2018 (cell) | 4.6439 | $2.36 \times 10^{-6}$ | 1.5076 | $3.18 \times 10^{-6}$ | Small |
| A/Victoria/653/2017 (cell) | 12.8883 | $1.51 \times 10^{-6}$ | 0.9455 | $2.20 \times 10^{-6}$ | Small |
| A/Victoria/653/2017 (egg) | 7.6359 | $1.93 \times 10^{-5}$ | 4.2875 | $2.33 \times 10^{-5}$ | Small |
| A/Switzerland/8060/2017(cell) | 7.1822 | $1.40 \times 10^{-7}$ | 1.7357 | $1.89 \times 10^{-7}$ | Small |
| A/Switzerland/8060/2017 (egg) | 5.3848 | $2.76 \times 10^{-6}$ | 1.5001 | $3.82 \times 10^{-6}$ | Small |

The magnitude of virus neutralization (approximately 10^0^ to 10^2^) and virus saturation (approximately 10^−1^ to 10^2^) remained relatively consistent among all 327 reactions with different virus-serum pairs (Fig. 2d and 2f). However, the antibody consumption rate formed a bimodal distribution, with the majority of reactions distributed at 10^−7^ to 10^−5^ (small antibody consumption rate) and small number reactions from 10^−1^ to 10^1^ (large antibody consumption rate) (Fig. 2c). A large antibody consumption rate would indicate a serum containing a greater proportion of weakly neutralising antibodies where several antibodies are required for effective neutralisation [6], whereas a low antibody consumption rate will like lead to rapid viral clearances due to rapid production of such antibodies suggests fewer antibody molecules required for neutralization [5]. A comparison of genetic distances between test and reference viruses showed that the categories by antibody consumption was independent of HA amino acid distance (Fig. 2d-g). A Pearson correlation analysis showed a positive correlation between antibody consumption*φ*) and antibody saturation (*η*) (correlation coefficient, 0.7601; Fig. 2d-g; Section 2.2, Supplementary material) and between virus neutralization (*α*) and antibody consumption (φ) (correlation coefficient, 0.7770) showing a strong relationship between virus neutralization by antibody binding and its consumption.

In some assays we observed that the virus with antisera grow better than controls without antisera (Table S2) by comparing focus numbers in well with antisera and that without antisera. We hypothesize that this may be due to the presence of virus growth factors or non-neutralising antibodies in the ferret antiserum, hence, to estimate virus neutralization parameter from FRA we used two approaches: most diluted antibody (qualified 327 datasets) and cell control as total viral titre (qualified 63 datasets) viral titre with the as total viral titre. However, regardless of selection of total titre, the magnitude and category of virus neutralization parameters remain consistent (Section 2.2, *Supplementary material*).

To obtain replication kinetics of influenza virus we fit a logistic growth model to previously published growth kinetics of H1N1pdm09 and H7N9 virus on A549 human lung carcinoma cells in the absence of antibodies (Simon et al. 2016 [33], Section 1.1, *Supplementary material*). Strikingly, the magnitudes of virus replication (~10^0^), virus degradation (~10^0^) and natural saturation (~10^−3^) were consistent for H1N1pdm09 and H7N9 virus are consistent (Section 1.1, *Supplementary material*), and since all subtypes have a similar function it is reasonable to assume that replication kinetics on A549 cells may follow a similar behaviour.

### Antibody-induced bistable viral kinetics

Integration of virus replication kinetics with the neutralization kinetics from FRA datasets for different magnitudes of antibody concentration (small and large, System 3), our results suggests that both virus neutralization kinetics with small and large antibody consumption lead to bistable viral kinetics (Figs. 3 and 4), but their behaviours are subtly different. For a small antibody consumption, exhibited by a majority of FRA data, we approximated the antibody concentration during the 144-hour incubation period as the initial antibody concentration (Fig. S9), and find that virus and antibody can co-exist at the end of the experiment (Fig. 3a-d, and S9). Further, the maximum capacity of viral titre decreases with increase of antibody concentration (Fig. 3e). On the other hand, for the few reactions that showed a large antibody consumption, viral survival corresponds to depletion of antibody, while viral eradication coincides with antibody remaining (Fig. 4 and Fig. S10). Moreover, both virus titres converged to the same maximum capacity if virus survives in presence of antibody (Fig. 4b, c and e).

**Fig. 3.**
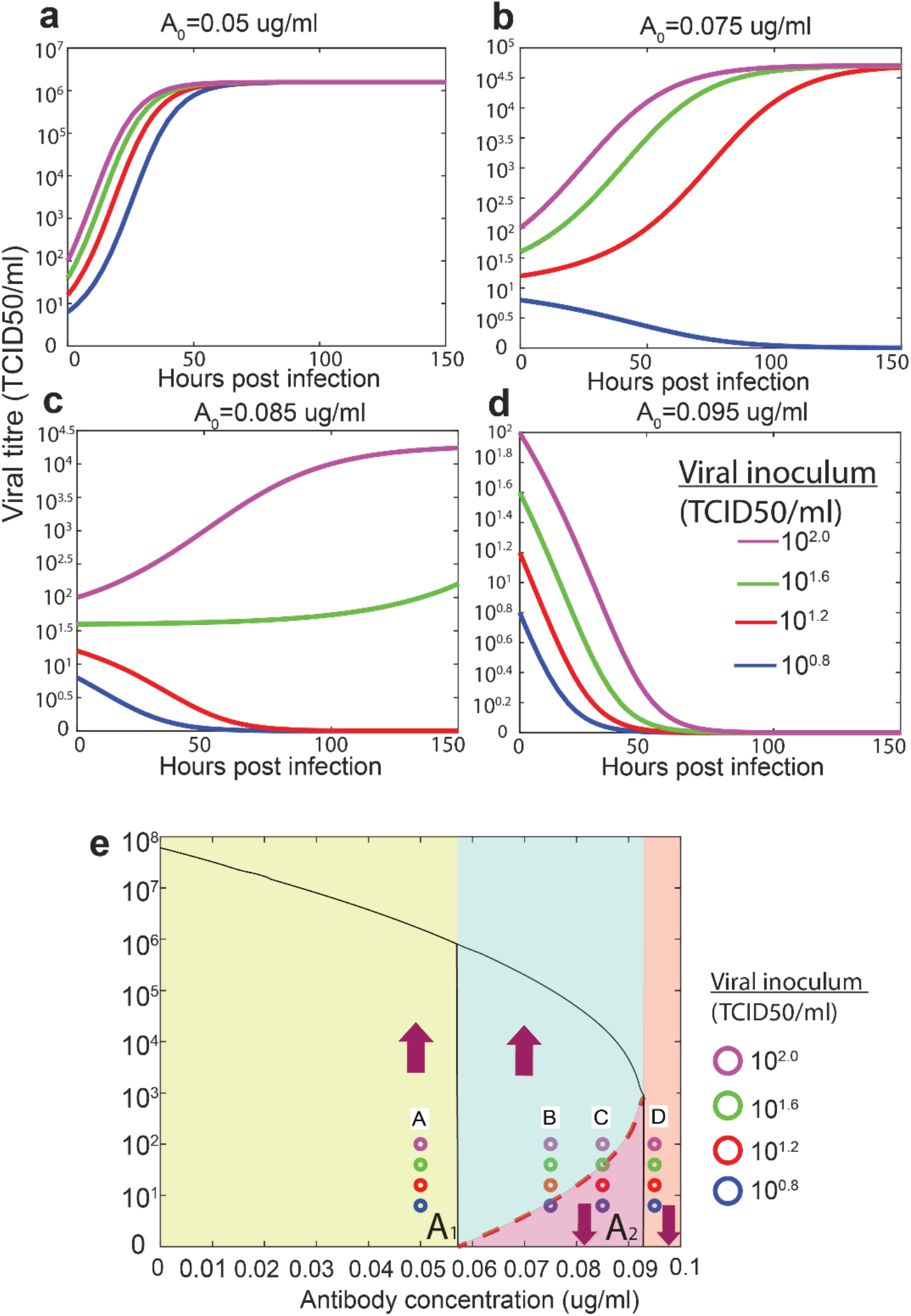
Simulated kinetics of H7N9 virus with different combinations of inoculum sizes and antibody concentrations with low antibody consumption. Bifurcation diagram e) showing viral titre as a function of antibody concentration. Viral inoculum threshold increases with increase of antibody concentration (red dashed curve). The maximal capacity of viral titre decreases with increases of antibody concentration (black solid curve). Purple arrows represent any viral inoculum size. A) When antibody concentration is between 0 and A_1_ virus with any viral inoculum survive. B) Virus with any inoculum size is inhibited when antibody concentration is greater than A_2_. C) and d) when antibody concentration is between A_1_ and A_2_ virus survives if viral inoculum is above dashed red curve and is inhibited if viral inoculum is below dashed red curve. Purple, green, red and blue lines and circles represent viral kinetics with viral inoculum 10^0-8^, 10^1-2^, 10^1-6^ and 10^2-0^ TCID50/ml in a–d, also shown as virus inoculum in e.

**Fig. 4.**
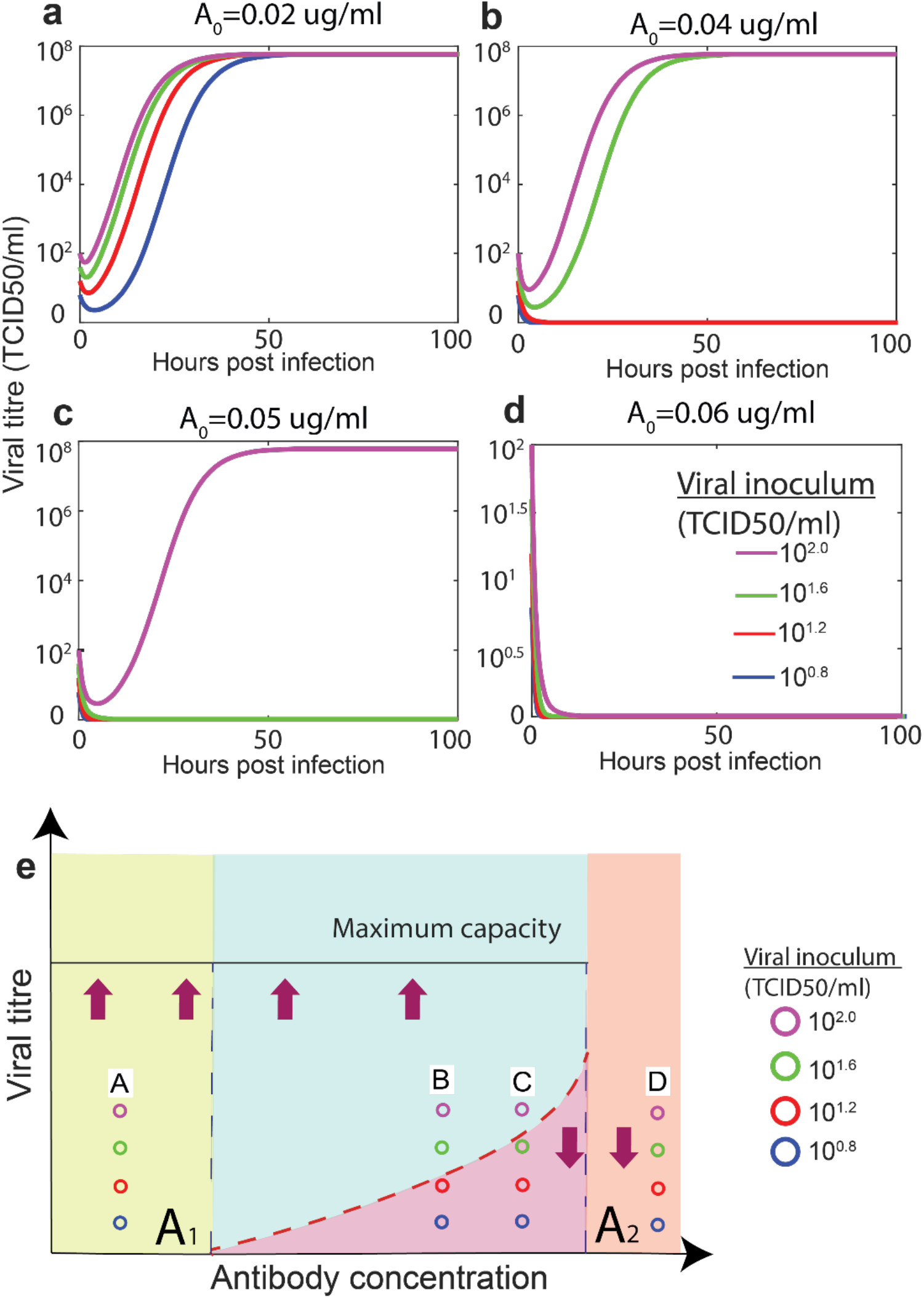
Simulated kinetics of H7N9 virus with different combinations of inoculum sizes and antibody concentrations with large antibody consumption. Viral survival corresponds to antibody depletion and viral eradication coincides with antibody existence. Bifurcation diagram e) showing viral titre as a function of initial antibody concentration (schematic diagram). Viral inoculum threshold increases with increase of antibody concentration (dashed red line). Maximum capacity of viral titre remains constant if virus survives (black solid line). Purple arrows represent any viral inoculum size. A) When antibody concentration is between 0 and A_1_, virus with any viral inoculum survive. B) and c) When antibody concentration is between A_1_ and A_2_, viral kinetics survives if viral inoculum is above dashed red curve and is inhibited if viral inoculum is below dashed red curve. D) Virus with any inoculum size is inhibited when antibody concentration is greater than A_2_. Purple, green, red and blue lines and circles represent viral kinetics with viral inoculum 10^0.8^, 10^1.2^, 10^1.6^ and 10^2.0^ TCID50/ml in a-d, also shown as virus inoculum in e.

In simulations of virus replication kinetics with saturated virus neutralization and small antibody consumption (Table S19), we found the existence of two thresholds *A*_1_ and *A*_2_ that divide the antibody concentration interval into three regimes, shown as bifurcation diagram (Fig. 3e). In different antibody concentration intervals, viral kinetics exhibits different dynamical behaviours. Virus kinetics exhibit bistability at the antibody concentration interval between *A*_1_ and *A*_2_, where small viral inocula are inhibited, and large viral inocula survive under the same antibody concentration (Fig. 3b and c). The viral inoculum threshold (above which the virus survives) increases with increase of antibody concentration (the red dashed curve in Fig. 3e). For example, at low antibody concentration *A* = 0.075 *ug/ml*, virus with inoculum 10^1.2^*TCID* 50/*ml*, 10^1.6^*TCID* 50/*ml* and 10^2.0^*TCID* 50/*ml* survive (Fig. 3b and e), whereas at high antibody concentration *A* = 0.085*ug/ml*, virus with high inoculum 10^0.6^*TCID* 50/*ml* and 10^2.0^*TCID* 50/*ml* survives (Fig. 3c and e). At antibody concentration less than the threshold *A*_1_, the virus survives independent of inoculum (Fig. 3a and e), and at antibody concentration higher than the threshold *A*_2_, the virus is inhibited independent of inoculum size (Fig. 3d and e).

Similarly, in simulations of virus kinetics with saturated virus neutralization and large antibody consumption (Table S18), a similar bistable behaviour was also observed (Table S18) (Fig. 4). With initial antibody concentration *A*_0_ = 0.04*ug/ml*, virus with inoculum 10^1.6^*TCID* 50/*ml* and 10^2.0^*TCID* 50/*ml* survives, while antibody is depleted (Figs. 4b, e and S10), with initial antibody concentration *A*_0_ = 0.05 *ug/ml*, only virus with inoculum 10^2.0^*TCID* 50/*ml* survives (Figs. 4c,e and S10).

To establish whether bistability depends on model structure, we conducted sensitivity analysis by adding an eclipse phase to the model (Section 1.2., *Supplementary material*). The existence of bistability and its mechanism were unchanged by the eclipse phase (Section 3.1.2 and 3.1.3, *Supplementary material*). The existence and mechanism of bistability were consistent when either FRA measurements of neutralised virus counts were used (either control cell or viral titre with the most diluted antibody as total viral titre) (Section 3.2, 3.3-3.4, *Supplementary material*). Lastly, simulations of virus replication using model parameter values for both seasonal H1N1 and avian H7N9 virus showed antibody-induced bistable virus kinetics (Section 3, *Supplementary material*).

### Unsaturated virus neutralization always leads to monostable virus kinetics

We hypothesize that unsaturated virus neutralization, commonly used to quantify virus neutralization by antibody binding *in vivo* and *in vitro* [20, 21, 23, 24, 26], would only lead to monostable virus kinetics (i.e. variability of virus neutralization would only be caused by antigenic change). Reconsidering virus neutralization kinetics (System 4, Method), we again found two categories of antibody consumption within our FRA data, including small and large antibody consumption (Table S22). In simulations of virus growth kinetics with unsaturated neutralisation and a small antibody consumption, neutralizing antibody only lead to monostable viral kinetics (Fig. 5); viral survivability only relied on magnitude of antibody concentration, rather than viral inoculum size. For large antibody consumption, however, neutralizing antibody lead to antibody-induced bistable viral growth kinetics exist (Fig. 6), but its bistable antibody concentration interval is relatively small. Thereby, we conclude that unsaturated virus neutralization leads to monostable viral kinetics under most conditions (Section 2.2.4, *supplementary material*), because small antibody consumption leads to monostable virus kinetics and large antibody consumption leads to bistable virus kinetics with small bistable antibody concentration interval. As a result, whether an antibody can neutralise a virus depends only on antibody concentration.

**Fig. 5.**
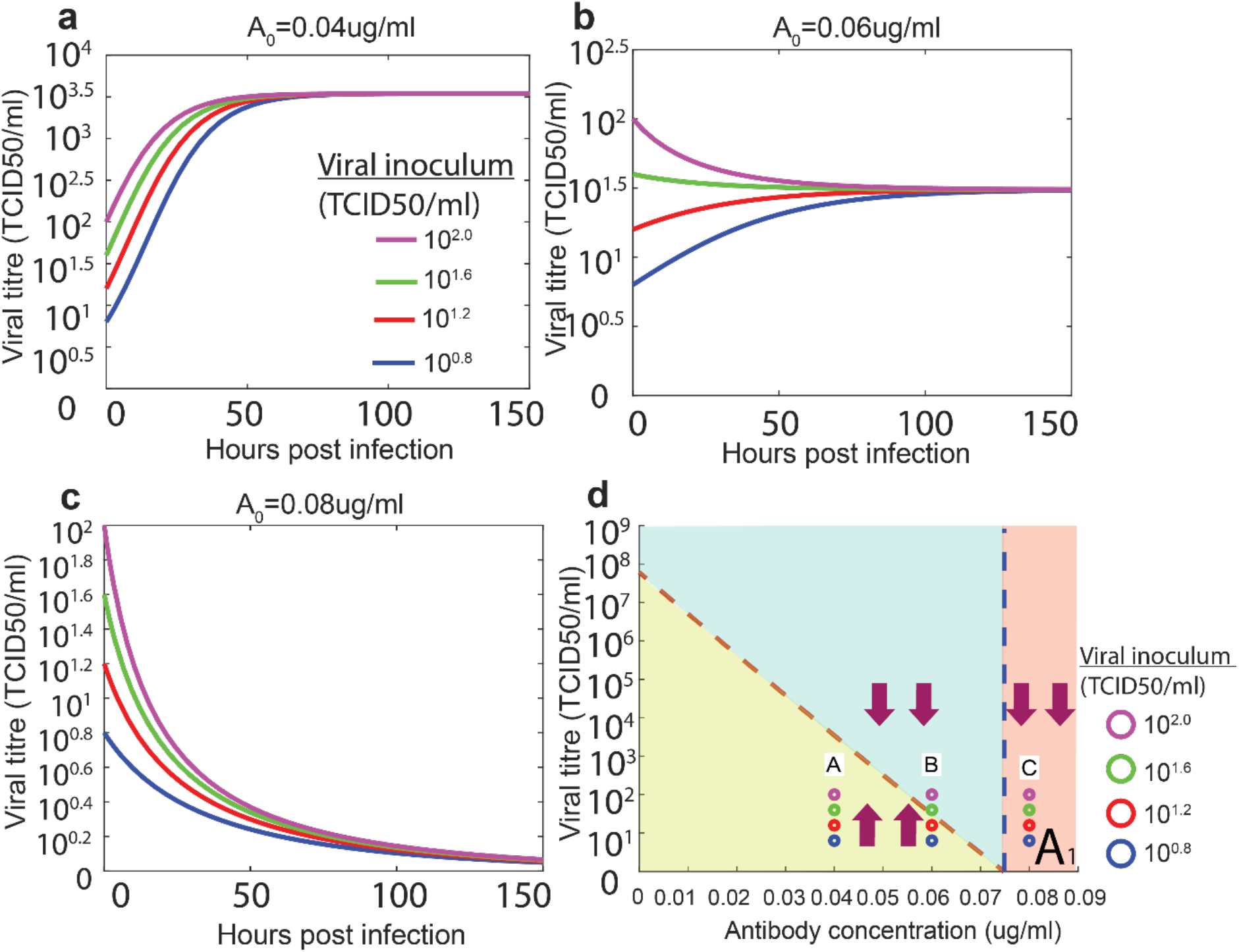
Simulated kinetics of A/H7N9 virus with different combinations of inoculum sizes and antibody concentrations with small antibody consumption. Bifurcation diagram d) showing viral titre as a function of initial antibody concentration. Maximal capacity of viral titre decreases with increase of antibody concentration (dashed red line). Purple arrows represent any viral inoculum size. A) and b) When antibody concentration is between 0 and A_1_, virus with any viral inoculum survive. C) Virus with any inoculum size is inhibited when antibody concentration is greater than A_1_. Purple, green, red, blue curve and circle represent viral kinetics with viral inoculum 10^0.8^, 10^1.2^, 10^1.6^ and 10^2.0^ TCID50/ml in (a-d).

**Fig. 6.**
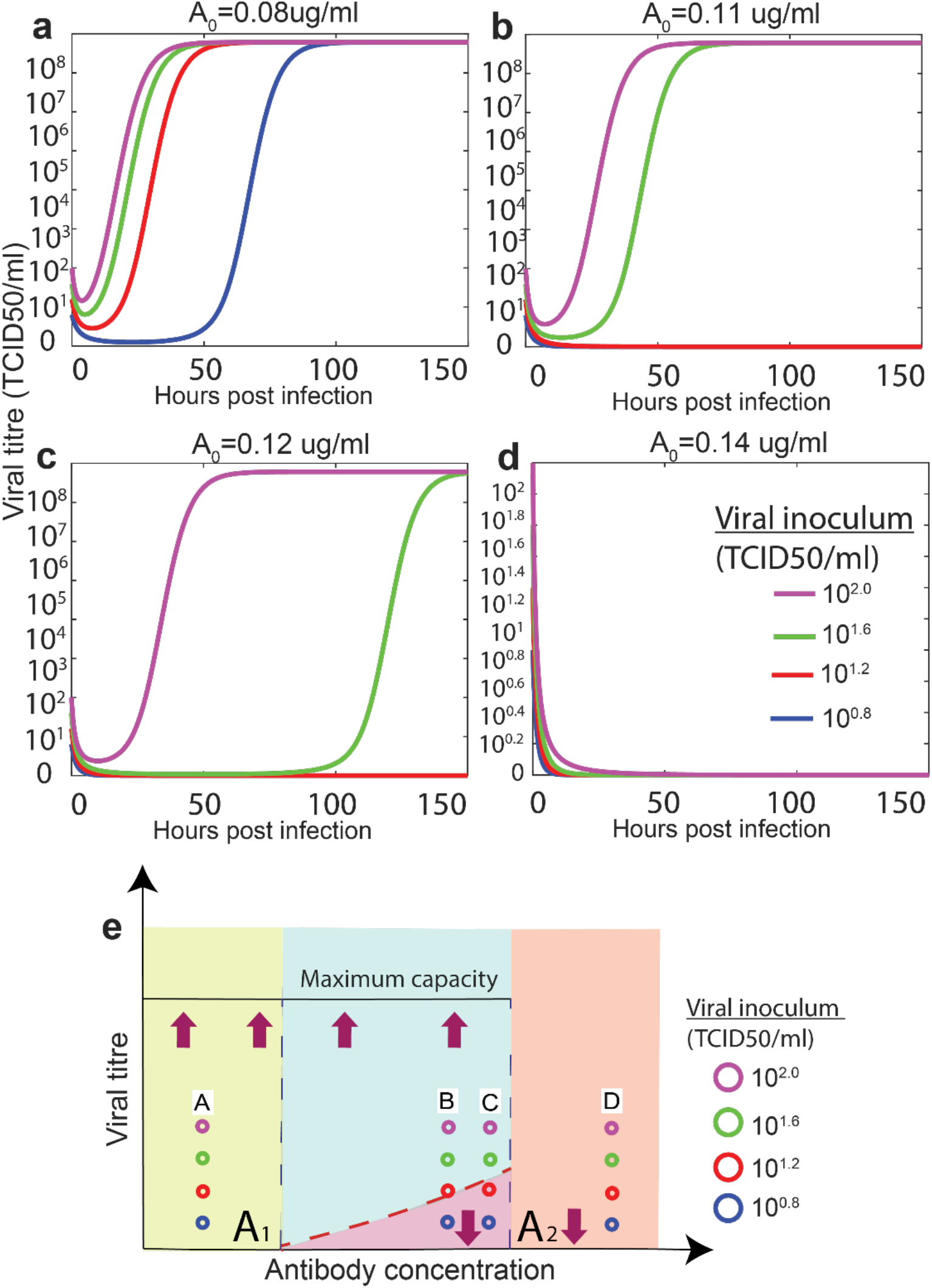
Simulated kinetics of A/H7N9 (virus with different combinations of inoculum sizes and antibody concentrations with large antibody consumption. Viral survival corresponds to antibody depletion and viral eradication coincides with antibody existence. Bifurcation diagram e) showing viral titre as a function of initial antibody concentration (schematic diagram). Viral inoculum threshold increases with increase of antibody concentration (dashed red line). Maximum capacity of viral titre remains constant if virus survives (black solid line). Purple arrows represent any viral inoculum size. A) When antibody concentration is between 0 and A_1_, virus with any viral inoculum survive. B) and c) When antibody concentration is between A_1_ and A_2_, viral kinetics survives if viral inoculum is above dashed red curve and is inhibited if viral inoculum is below dashed red curve. D) Virus with any inoculum size is inhibited when antibody concentration is greater than A_2_. Purple, green, red, blue curves and circle represent viral kinetics with viral inoculum 10^0.8^, 10^1.2^, 10^1.6^ and 10^2.0^ TCID50/ml in (a-d).

To summarize, we found that saturated virus neutralization can always lead to antibody-induced bistable viral kinetics (Section 3, *Supplementary material*). Unsaturated virus neutralization leads to monostable viral kinetics for small consumption (Section 4, *Supplementary material);* unsaturated virus neutralization leads to bistable viral kinetics, but its bistable antibody concentration interval is relatively small (Section 4, *Supplementary material*) (Table 3).

**Table 3.** Bistability/Monostability determined by virus neutralization, eclipse phase and antibody consumption

| Model | Virus neutralisation | Eclipse phase | Antibody consumption | Bistability or monostability |
| --- | --- | --- | --- | --- |
| 1 | Saturated | No | Small | Bistability |
| 2 | Saturated | No | large | Bistability |
| 3 | Saturated | Yes | small | Bistability |
| 4 | Saturated | Yes | large | Bistability |
| 5 | Unsaturated | No | small | Monostability |
| 6 | Unsaturated | No | large | Bistability, small antibody concentration interval |
| 7 | Unsaturated | Yes | small | Monostability |
| 8 | Unsaturated | Yes | large | Bistability, small antibody concentration interval |

## Discussion

Dissecting the neutralised viral titer estimated obtained for 49 A/H3N2 viruses circulating during 2014-2019 against eight reference antisera raised in ferrets, we identified that the antibody consumption rates of the reaction formed two distinct groups, with either a small or large antibody consumption rate and this correlated strongly with antibody saturation. The differences in antibody consumptions rates were not associated with influenza HA genetic distances, although our study included test and reference viruses with high genetic and antigenic similarity. Taken together, the variation in antibody consumption into distinct categories suggests that this may be due to interacting factors that affect the potency of the serum, such as binding avidity and affinity.

By integrating estimated viral replication and virus neutralization parameters, we illustrated that neutralising antibodies induce bistable viral kinetics through saturation of virus neutralisaiton and antibody consumption. Biologically, even for a well matched virus-antibody pair, large viral inocula survive and small viral inocula are inhibited at the same antibody concentration. This supports that escape from neutralization can result from innate interactions between well-matched virus strain and antibodies. Our results imply that even for the same virus-antibody pair, the elimination of virus depends not only on the antibody concentration but also virus inoculum size, highlighting their important roles in the establishment of a successful infection.

Our results also imply that antibody levels measured using HI assays, may be inadequate to measure protection as the HAI titres only show the effect of varying antibody concentration, but not the effect of varying the virus inoculum. On the other hand, analysis of FRA data using mathematical models enables us to understand both of these factors. During a vaccine selection process, viruse strains that induce antibodies with high virus neutralization and small antibody consumption would be favourable.

HI and FRA are designed to quantify virus replication kinetics of antisera arisen from test virus. Antibody-antisera mixture is incubated for 30 minutes for HAI and one hour for FRA, and infectious viral titre is decreasing during antibody-virus incubation. However, the result provided by short-time incubation assay may not reflect the whole-picture of virus neutralization kinetics, because virus titre *in vivo* changes with respect of time, for example logistics model due to limited susceptible cells and a bimodal growth kinetics due to interferon [22]. Combination between virus replication kinetics and saturated virus neutralization leads to variability of virus neutralization independent of antigenic changes, indicating that incorporation of virus replication kinetics is an urgent need for future assay to quantify virus replication kinetics.

A variety of proposed target cell-infected infected cell-virus (TIV) models of hepatitis B virus (HBV) and simian immunodeficiency virus (SIV) exhibit bistable viral kinetics, however bistability has been attributed to reversible binding of free antibody, which may be biologically unreasonable[23, 24]. Further, these models model virus neutralization as unsaturated, using proportional to product of antibody concentration and viral titre by law of mass action, which is unrealistic as discussed in the Introduction. To our best knowledge, the proposed model is the simplest model with realism leading to bistable switch between viral survival and eradication. By Occam’s razor principle, because saturated virus neutralization and antibody concentration is the simplest adequate model, it provides a main underlying mechanism to explain variability of virus neutralization.

A limitation of our viral replication kinetics model (in the absence of antibodies) is that it cannot reproduce observed viral titres when the inoculum is close to the maximum viral load. While A/H1N1pdm09 and A/H7N9 viruses fit our model, the seasonal A/H1N1 (sH1N1) and A/H5N1 data from the same study^35^ did not achieve a good fit (data not shown), possibly because the inoculum level is closer to the peak viral load for these data. However, since the viral inoculum in natural infection is considerably low, our proposed one-dimensional model should be adequate to capture viral kinetics of natural infection. For low inocula, we are confident in our model because for each virus, we fitted the model to data from two inocula simultaneously (0.01 PFU/cell and 3 PFU/cell), and produced good fits for both inocula.

We note that when we qualitatively analyse the long-term behaviour of our models, virus always survives and antibody is always depleted. However, within a realistic timeframe for experimental and natural infection (144 hours), bistability exists. Also, if the viral load is low after 144 hours, even though a deterministic model would predict it to rebound once antibodies are depleted, in reality stochastic effects would eradicate the virus before this could happen (data not shown).

Using viral replication parameters from one experiment and virus neutralization parameters from another experiment, we have predicted the existence of antibody-induced bistable viral kinetics. To identify the bistable antibody interval for a specific antibody-antigen pair, further work is required to experimentally validate this prediction in a single experimental system. Moreover, natural infection can occur with a mixture of virus genotypes (within-host genetic diversity), and the antibody response produced in response to natural infection is polyclonal, hence future modelling with a mixture of genotypes, and protection by polyclonal antibodies is required. Also, while viral inoculum size (initial viral load) is the major determinant of survival or death *in vitro, in vivo* survival of influenza is determined by factors beyond virus inoculum size, including the time of antibody production and other innate immunity functions. Thus, although antibody concentration and inoculum size are two seemingly important factors, they are not sole contributors for *in vivo* virus eradication. Also, our experimental system does not consider variations in growth kinetics in different sites (e.g. nasal vs lung). Further, in our study we used naïve ferret raised antisera where the primary response is known to be narrow, in contrast to humans who exhibit a complex immune history.

## Method

### Mathematical model

To establish quantitative relationship between neutralized viral titre and antibody concentration, we developed two models of (a) saturated virus neutralization (System 1, below) and (b) unsaturated virus neutralization (System 2, below).

For saturated virus neutralization, we describe the rate of change of viral titre and antibody concentration as 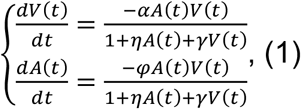 in FRA for one-hour incubation. *A*(*t*) represents antibody concentration with respect to time *t. α* represents the virus neutralization rate by antibody binding; *φ* represents antibody consumption rate by binding to virus; *η* controls the saturation in neutralization rate as antibody concentration increases; *γ* controls the saturation in neutralization rate as viral titre increases.

For unsaturated virus neutralization, we describe the rate of change of viral titre and antibody concentration as 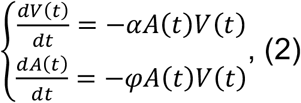, in FRA for one-hour incubation.

To investigate the role of antibody concentration and viral inoculum on viral kinetics we developed four models of viral kinetics (a) with saturated virus neutralization and antibody consumption (System 3, below); (b) with saturated virus neutralization, antibody consumption and with eclipse phase (System 5, Section 3.2., *Supplementary Material);* (c) with unsaturated virus neutralization and antibody consumption (System 4, below); and (d) unsaturated virus neutralization, antibody concentration and eclipse phase (System 6, *Supplementary Material*).

For viral kinetics with saturated virus neutralization and antibody consumption, we describe the rate of change of viral titre and antibody concentration as 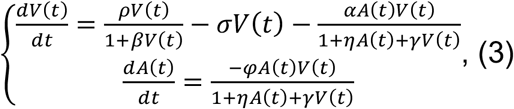, to simulate *in vitro* experiment, where *ρ* represents replication rate of influenza virus; *β* controls natural saturation of viral replication at high viral titer; *σ* represents degradation rate of influenza virus.

For viral kinetics with unsaturated virus neutralization and antibody consumption, we describe the rate of change of viral titre and antibody concentration as 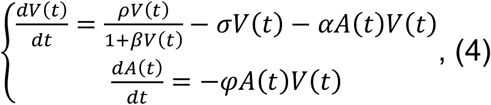, to simulate in vitro experiment.

To understand the antigenic relationships among tested H3N2 viruses, we compared the amino acid distance between their hemagglutinin genes (HA) using MEGA X (https://www.megasoftware.net/) [34]. The HA genes of newly generated and reference strains are available in the Global Initiative on Sharing All Influenza Data (GISAID) database (https://www.gisaid.org/) [35]. Sequence accession numbers and laboratories generating the sequence data are provided in the Supplementary Materials.

### Infection assay

Virus replication parameters were obtained from single cycle (SC) and multiple cycle (MC) infection assays performed by Simon *et al*. [33]. A549 human lung carcinoma cells were infected with influenza A/H1N1pdm09 (A/Mexico/INDRE4487/2009) and A/H7N9 (A/Anhui/1/2013). Both a high viral multiplicity of infection (MOI) (3 PFU/cell) and a low MOI (0.01 PFU/cell) were used. 0.5 mL of the cell supernatant was harvested and frozen at 13 intervals for SC assay that lasted 18 hours (0, 1, 2, 3.5, 4.6, 5.6, 7.1, 8.6, 10, 11, 12, 15.5 and 18 hours) whereas 11 intervals (0, 3.1, 18.6, 28.5, 42.9, 53.3, 66.5, 77.8, 91.6, 99 and 147) were sampled for MC infection assays that lasted ~150 hours. The frozen samples were thawed and titrated by median tissue culture infectious dose (TCID50) by Simon *et al*^35^.

### Focus Reduction assay

To determine parameters of virus neutralization we utilized a focus reduction assay (FRA) performed against 49 H3N2 viruses using post-infection ferret antisera raised against a panel of representative H3N2 viruses. Serial dilutions (80 to 10240) of ferret antisera were incubated for one hour with virus and diluted to 1000 FFU/well. 100ul of the virus-sera mixture was then applied to confluent MDCK-SIAT cells and incubated for 18-20 hours at 35°C in 5%CO2. FRA was performed for each virus individually with each ferret antisera raised against a representative set of H3N2 viruses. In the main-text, we describe the dynamics using A/Canberra/40/2019. The virus-sera mixture was then added to confluent MDCK-SIAT1 cell lines, allowing the measurement of antibodies required to neutralise virus through reduction of plaques during one-hour incubation. Following overnight incubation, focus forming units (FFU) were quantified by immunostaining using an anti-nucleoprotein monoclonal antibody and subsequent detection using an HRP-conjugated secondary antibody (BioRad, USA) and TrueBlue substrate (KPL Biosciences). The number of FFU per well was quantified from plate images using an Immunospot analyser and Biospot software (CTL Immunospot, USA).

### Parameter estimation

Variation in viral replication parameters was estimated using the combined sum of squared error (combined SSE) across the single-cycle and multi-cycle experiments. Combined SSE is 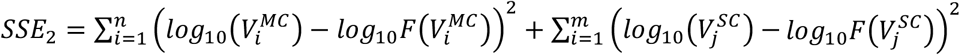, where 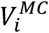 and 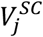 represents experimental viral titer at time *i* and *j. F*(*V_i_*) and *f*(*V_j_*) represent estimated viral titer at time *i* and *j* for SC or MC infection assay data. Initial guesses used for parameter estimation are *ρ*_0_ = 1, *β*_0_ = 0.1, *σ*_0_ = 1 and *τ* = 0.1.^#^

For the estimation of virus neutralization parameters, we obtained neutralized influenza virus in one-hour of incubation by using the formula neutralized virus = total virus – survived virus, and selected column with all positive values. Then we calculated neutralized viral titer by the formula, *Viraltiter = FN × DR × SV FFU/ml* where *FN* represents focus number in each well, *DR* represents dilution rate, *SV* represents sample volume and *FFU* represents focus formation assay. We used the 7 virus serial dilution data points (80 – 5120) of the FRA, where *A_i_* represents diluted antibody concentration and *V_i_* represents neutralized virus titer. We defined the sum-of-squares error (SSE) as 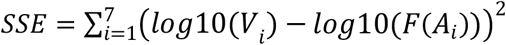, where *V_i_* and *F*(*A_i_*) represents experimental and theoretical viral titers with respect to the i^th^ diluted antibody concentration. *F*(*A_i_*) is the integral of 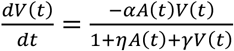 from 0 to 1.

### Qualitative analysis of Mathematical models

By Bendixon-Poincare theorem [28, 30, 36], we show non-existence of closed orbits in model systems of viral kinetics with antibody consumption, with and without eclipse phase (System 3 and 4, respectively). The existence of equilibria and its stability for systems with no antibody consumption (the first equation of System 3 and 4) is provided by reference^24^. The introduction of eclipse phase in System 5 and System 6 in supplementary material does not change the stability of equilibria provided by System 3 and System 4, respectively, as shown by Rouche’s Theorem [37] and the continuity of eclipse phase*τ*.

## Acknowledgements

We acknowledge the technical support and advice of Dr. Brendan Russ, Dr. Claerwen Jones, Dr. Jasmine Li and Prof. Zhongfang Wang. S.X and D.V. are supported by contract HHSN272201400006C from the National Institute of Allergy and Infectious Diseases, National Institutes of Health, U.S. Department of Health and Human Services, USA; A.W.C.Y. is supported by a Wellcome Trust Collaborative Award (200187/Z/15/Z); C.M.D is supported by an Australian National Health and Medical Research Council Early Career Fellowship (1113269). The Melbourne WHO Collaborating Centre for Reference and Research on Influenza is supported by the Australian Government Department of Health.

