## Supplementary document for "Saturation of influenza virus neutralization and antibody consumption can both lead to bistable growth kinetics"

This pdf file includes:

**Supplementary text**

**Figures S1 to S36**

**Tables S1 to S36**

#### Supplementary Information text

#### Appendix - Table of Contents

1. Virus replication kinetics of influenza virus (H7N9 and H1N1pdm09)
  1.1. Virus replication kinetics without eclipse phase- Ordinary differential equation (ODE) model
  1.2. Virus replication kinetics with eclipse phase- Delayed differential equation (DDE) model

2. Virus neutralization parameter estimation from Focus Reduction Assay data
  2.1. Calculation of neutralized virus titre
    *2.1.1. Neutralized viral titre estimated from cell control as total viral titre*
    *2.1.2. Neutralized viral titre estimated from viral titre with the most diluted antisera as viral titre*
  2.2. Estimation of virus neutralization parameters
    *2.2.1. Estimation of virus neutralization parameters – FFU scale (cell control as total viral titre)*
    *2.2.2. Estimation of virus neutralization parameters – TCID50 scale (cell control as total viral titre)*
    *2.2.3. Neutralization parameters of A/H3N2 viruses – FFU scale (cell control as total viral titre)*
    *2.2.4. Estimation of virus neutralization parameters – FFU scale (most diluted antisera as total viral titre)*
    *2.2.5. Estimation of virus neutralization parameters – TCID50 scale (most diluted antisera as total viral titre)*
    *2.2.6. Neutralization parameters of A/H3N2 viruses – FFU scale (most diluted antisera as total viral titre)*

3. Bistable kinetics of influenza virus in the presence of neutralizing antibody
  3.1. Virus neutralization parameter estimated from viral titre with the most diluted antisera as total viral titre
    *3.1.1. Antibody concentration kinetics of A/H7N9 viral replication parameter in presence of antibody*
    *3.1.2. Bistable kinetics of A/H7N9 with small consumption rate and eclipse phase*
    *3.1.3. Bistable kinetics of A/H7N9 with large consumption rate and eclipse phase*
  3.2. Virus neutralization parameter estimated from cell control as total viral titre
    *3.2.1. Bistable kinetics of A/H1N1pdm09 with small and large antibody consumption rate*
    *3.2.2. Bistable kinetics of A/H7N9 with small and large antibody consumption rate*
  3.3. Bistable kinetics of A/H1N1pdm09 with eclipse phase and small antibody consumption or large antibody consumption
  3.4. Bistable kinetics of A/H7N9 with eclipse phase and small antibody consumption or large antibody consumption

4. Viral kinetics and antibody concentration kinetics with unsaturated virus neutralization
4.1. Virus neutralization parameter estimated from viral titre with the most diluted antisera as total viral titre
4.1.1. *Antibody concentration kinetics of A/H7N9 viral replication in presence of antibody*
4.2. Virus neutralization parameter estimated from viral titre with cell control as total viral titre
4.2.1. *Monostable kinetics of A/H1N1pdm09 with small antibody consumption rate*
4.2.2. *Bistable kinetics of A/H1N1pdm09 with large antibody consumption rate*

5. GISAID accession numbers and acknowledgements to generating labs

### 1. Virus replication kinetics of influenza virus (H7N9 and H1N1pdm09)

In this section, we describe the estimation of virus replication parameter from both single-cycle (SC) and multiple-cycle (MC) infection assay of A/H1N1pdm09 and avian influenza A/H7N9 viruses with ordinary differential equation (ODE) model (Section 1.1) and delay differential equation (DDE) model (Section 1.2). We believe that stochastic effects may not play a decisive role supporting the usage of ODE and DDE models, as viral titres are considerably large during both clinical and experimental infection (in the range of $10^6 \sim 10^8\ TCID50/ml$)[1-4], although stochasticity may affect the steady state reached from a given initial condition, such as if early extinction of the virus occurs.

#### 1.1. Virus replication kinetics without eclipse phase- Ordinary differential equation (ODE) model

Since, in the absence of antibodies influenza replication *in vitro* and *in vivo* is solely limited by the availability of susceptible cells [1, 2, 5], we describe the change of virus titre through natural virus replication and degradation in limited number of susceptible cells as the one-dimensional ODE model, $\frac{dV(t)}{dt} = \frac{\rho V(t)}{1+\beta V(t)} - \sigma V(t)$, (1). This system is illustrated by the green terms in Fig. S1. $V(t)$ represents viral titre with respect to time $t$. Parameters $\rho$ represents replication rate of influenza virus; $\beta$ controls natural saturation of viral replication at high viral titre; $\sigma$ represents degradation rate of influenza virus.

The SC assay was performed with infection at high viral multiplicity of infection (MOI), 3 plaque forming unit (PFU)/cell, such that 95% of the cells were infected simultaneously, whereas the MC assay was performed with infection at low MOI (0.01 PFU/cell) such that only a small population of cells were infected and new viral progeny are continuously generated. Thus, SC and MC assays provide viral replication kinetics with different inoculum sizes.

We obtained the fitted curves with parameters through combined sum-of-squares error (combined SSE); defined as the combined SSE of SC and MC infection assay data, $SSE = \sum_{i=1}^{n}\left(log_{10}\left(V_i^{MC}\right) - log_{10}F\left(V_i^{MC}\right)\right)^2 + \sum_{i=1}^{m}\left(log_{10}\left(V_j^{SC}\right) - log_{10}F\left(V_j^{SC}\right)\right)^2$, (2) where $V_i^{MC}$ and $V_j^{SC}$ represents experimental viral titre at time $i$ and $j$. $F(V_i)$ and $F(V_j)$ represent estimated viral titre at time $i$ and $j$ for SC or MC infection assay data. We fit

System 1 to single-cycle (SC) and multiple-cycle (MC) infection assay of A/H1N1pdm09 and avian influenza A/H7N9 viruses, performed by Simon *et al*.[2], using the combined sum of squared error (combined SSE) (System 2).

Regardless of inoculum size, the kinetics of both A/H1N1pdm09 and A/H7N9 converged to the same maximal capacity (Fig. S2). Virus replication parameters obtained through combined SSE can fit SC and MC infection assay data of both viruses (Table S1), and is used to examine the role of virus neutralization in viral kinetics in main text and supplementary material.

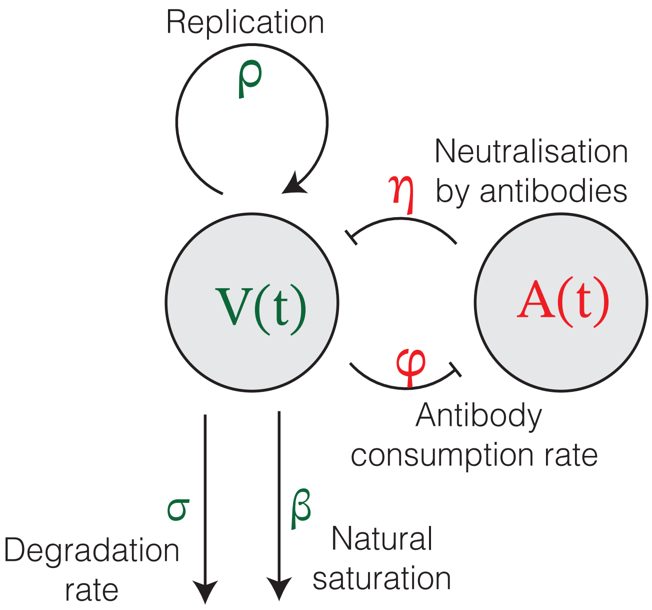


**Figure S1.** Flowchart of influenza kinetics in the presence of neutralizing antibodies. Viral kinetics is determined by limited viral growth, saturated virus neutralization and saturated antibody consumption.

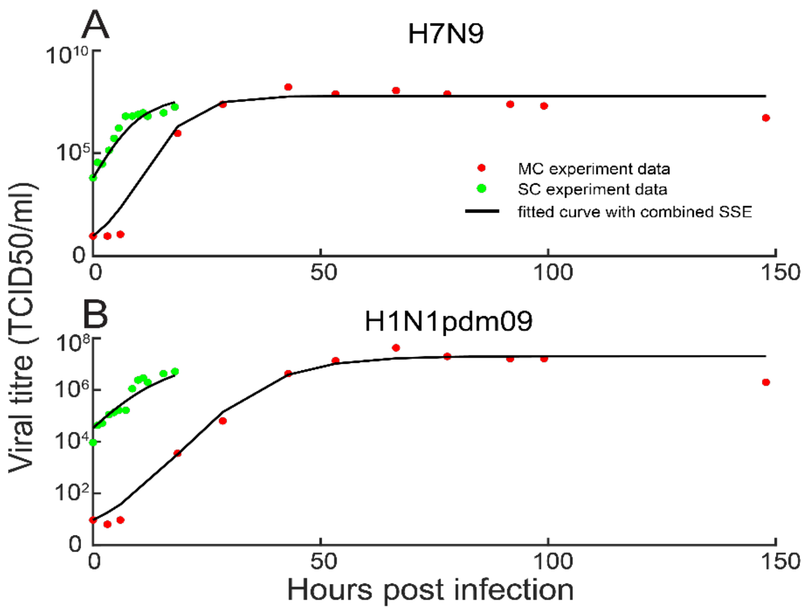


**Figure S2.** Viral replication kinetics of influenza virus on A549 human lung carcinoma cells. Regardless of viral inoculum sizes, used in single cycle (SC, green) and multi cycle (MC, red) infection assays, both viruses H7N9 (A) and H1N1pdm09 (B) converge to the same maximal capacity. Virus replication parameters were obtained through combined SSE, simulated for 18 hours for the SC assay and 144 hours for MC infection assay. Black curve represents simulated viral replication curve.

**Table S1.** Estimated parameters of viral replication kinetics (TCID50 scale)

| Strain* | Natural birth rate, $\rho$, (TCID50/ ml/hour) | Saturation growth parameter, $\beta$, (hour/TCID50/ml) | Natural death rate, $\sigma$, (TCID50/ ml/hour) | Combined SSE |
|---|---|---|---|---|
| H1N1pdm09* | 1.5555 | 0.0093 | 1.4565 | 3.0821 |
| H7N9* | 4.4725 | 0.0056 | 4.2857 | 5.552 |

*$\rho_0 = 1, \beta_0 = 0.1, \sigma_0 = 1$

#### 1.2. Virus replication kinetics with eclipse phase- Delayed differential equation (DDE) model

We extended the one-dimensional delayed differential equation (DDE) described in Section 1.1 to model the replication kinetics of influenza in cell culture, including an eclipse phase $\tau$ (Fig. S3). Rate of change of influenza with an eclipse phase in cell culture is defined as $\frac{dV(t)}{dt} = \frac{\rho V(t-\tau)}{1+\beta V(t)} - \sigma V(t)$, (3).

For SC infection assay data, virus kinetics with viral replication parameters through combined SSE is simulated for 18 hours (Fig. S4); whereas MC is simulated for 144 hours (Fig. S4). Virus replication parameters obtained through combined SSE provided good fit for both SC and MC data (Table S2). Moreover, virus replication parameters obtained for both H7N9 and H1N1pdm09 through combined SSE can fit both SC and MC data, showing that regardless of viral inoculum sizes, kinetics of both virus subtypes converge to its same maximal capacity.

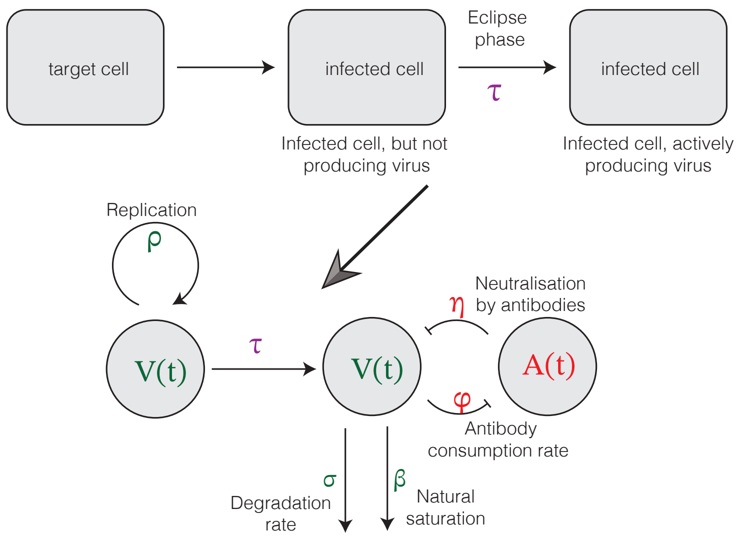


**Figure S3**. Flowchart of replication kinetics with eclipse phase of H7N9 and H1N1pdm09 virus on A549 cells

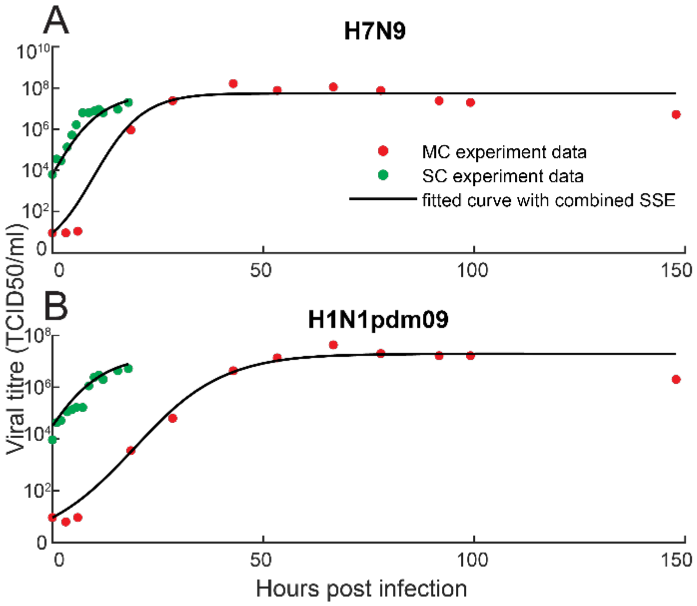


**Figure S4.** Virus replication kinetics (H7N9 and H1N1pdm09) on A549 human lung carcinoma cells. Regardless of viral inoculum size, influenza virus kinetics converges to same maximal capacity, A) H7N9 and B) H1N1pdm09. For both H7N9 and H1N1pdm09, virus replication parameter obtained through combined SSE can fit MC (red points) and SC (green point) infection assay data. For SC infection assay, virus kinetics with parameter through combined SSE is simulated for 18 hours; for MC infection assay, virus kinetics with parameter through combined SSE is simulated for 144 hours. Black curve represents simulated virus replication curve.

**Table S2.** Estimated parameters of virus replication kinetics (TCID50 scale)

| Strain* | Natural birth rate, $\rho$, (TCID50/ ml/hour) | Saturation growth parameter, $\beta$, (hour/TCID50/ml) | Natural death rate, $\sigma$, (TCID50/ml/hour) | Eclipse phase, $\tau$, (hours) |
|---|---|---|---|---|
| H1N1pdm09(A) | 0.9149 | 0.0193 | 0.8021 | 0.1235 |
| H7N9 | 0.8896 | 0.0468 | 0.6531 | 0.1477 |

*Initial parameters are $\rho_0 = 1$, $\beta_0 = 0.1$, $\sigma_0 = 1$ and $\tau = 0.1$.

### 2. Virus neutralization parameter estimation from Focus Reduction Assay

Below, we describe the estimation of virus neutralization from raw Focus Reduction Assay (FRA) data to obtain neutralized virus titre in two approaches, including cell control as total viral titre (Section 2.1.1) and viral titre with the most diluted antisera (Section 2.1.2). Next, we propose three models to fit neutralized virus titre on both scales and estimate the neutralization parameters, including focus forming units (FFU) (Section 2.2.1 and Section 2.2.3) and median tissue culture infectious dose (TCID50) (Section 2.2.2 and Section 2.2.4).

#### 2.1. Calculation of neutralized virus titre

The number of neutralised viruses in one-hour incubation were estimated from raw FRA (Table S3) using A/Brisbane/37/2017. The formula, neutralized virus = total virus - survived virus (Table S4, S5). Columns 1 to 10 in Table S3 represents post-infection ferret antisera raised against ten reference H3N2 viruses, including cell-grown A/Hong Kong/4801/2014, cell and egg grown A/Newcastle/82/2018, cell-grown A/Sydney/22/2018, cell and egg grown A/Victoria/653/2017, cell and egg grown A/Switzerland/8060/2017. Because viral titre with antisera is higher than virus without antisera for some virus-antisera combinations (Table S3), we use two approaches to estimate virus neutralization parameter, including cell control used as total viral titre and viral titre with the most diluted antisera used as total viral titre. Then, we use each column as survived virus with diluted antibody concentrations. Regardless of selection of total viral titre, the magnitude and category of virus neutralization parameter remains consistent. Then, we will show cell control as total viral titre (Section 2.1.1) first and viral titre with the most diluted antisera as total viral titre next (Section 2.1.2).

**Table S3**. Raw Focus Reduction Assay data generated using A/Brisbane/37/2017.

| Serum | Antisera arisen from reference virus (Focus number) | Cell control |
|---|---|---|

| Dilution | A/Hong Kong/4801/2014 (Egg grown) | A/Newcastle/82/2018 (cell grown) | A/Newcastle/82/2018 (egg grown) | A/Sydney/22/2018 (cell grown) | A/Victoria/653/2017 (cell grown) | A/Victoria/653/2017 (egg grown) | A/Switzerland/8060/2017(cell grown) | A/Switzerland/8060/2017 (egg grown) | |
|---|---|---|---|---|---|---|---|---|---|
| 80 | 72 | 97 | 81 | 49 | 58 | 43 | 15 | 109 | 3535 |
| 160 | 298 | 35 | 38 | 67 | 21 | 8 | 17 | 20 | 2422 |
| 320 | 1858 | 98 | 480 | 16 | 16 | 11 | 84 | 8 | 2104 |
| 640 | 2921 | 577 | 1889 | 17 | 29 | 128 | 328 | 16 | 2621 |
| 1280 | 3367 | 1701 | 2307 | 383 | 62 | 326 | 1105 | 97 | 2633 |
| 2560 | 4021 | 3424 | 3443 | 2340 | 64 | 1364 | 1789 | 692 | 2388 |
| 5120 | 4010 | 2496 | 2670 | 2568 | 270 | 1659 | 2630 | 1302 | 3043 |
| 10240 | 3726 | 2944 | 3436 | 2462 | 1574 | 1971 | 2844 | 2037 | 1440 |
| | | | | | | | | [*]AVE | 2678 |
| | | | | | | | | [+]SD | 620.2676 |

[*]AVE and [+]SD represents average and standard deviation.

##### *2.1.1. Neutralized viral titre estimated from cell control as total viral titre*

Below, we estimate neutralized viral titre from cell control and use averaged focus number in cell control as total viral titre. The neutralised viral titre in each well (Table S5) is obtained by substituting neutralized focus number in each well (Table S4) into the formula, $Viral\ titre = FN \times 20 \times 10^{3.5}\ FFU/ml$, where viral dilution factor is $10^{-3.5}$ and 50ul diluted virus in each well, $FN$ represents focus number in each well with one focus formation unit (FFU) representing one infectious influenza virus. Similarly, we obtain viral titre of total virus: $2678 \times 20 \times 10^{3.5} = 1.6937 \times 10^{8}$ FFU/ml.

**Table S4**. Neutralized foci number in each well (cell control as total viral titre)

| Serum Dilution | A/Sydney/22/2018 (cell grown) | A/Victoria/653/2017 (cell grown) | A/Victoria/653/2017 (egg grown) | A/Switzerland/8060/2017 (egg grown) |
|---|---|---|---|---|
| 80 | 2629 | 2620 | 2635 | 2569 |
| 160 | 2611 | 2657 | 2670 | 2658 |
| 320 | 2662 | 2662 | 2667 | 2670 |
| 640 | 2661 | 2649 | 2550 | 2662 |

| | | | | |
|---|---|---|---|---|
| 1280 | 2295 | 2616 | 2352 | 2581 |
| 2560 | 248 | 2614 | 1314 | 1986 |
| 5120 | 110 | 2408 | 1019 | 1376 |
| 10240 | 216 | 1104 | 707 | 641 |

**Table S5**. Neutralized viral titre in each well (FFU/ml, cell control as total viral titre)

| Serum Dilution | A/Sydney/22/2018 (cell grown) | A/Victoria/653/2017 (cell grown) | A/Victoria/653/2017 (egg grown) | A/Switzerland/8060/2017 (egg grown) |
|---|---|---|---|---|
| 80 | $1.6627 \times 10^8$ | $1.657 \times 10^8$ | $1.6665 \times 10^8$ | $1.6247 \times 10^8$ |
| 160 | $1.6513 \times 10^8$ | $1.680 \times 10^8$ | $1.6886 \times 10^8$ | $1.6801 \times 10^8$ |
| 320 | $1.6835 \times 10^8$ | $1.683 \times 10^8$ | $1.6867 \times 10^8$ | $1.6886 \times 10^8$ |
| 640 | $1.6829 \times 10^8$ | $1.675 \times 10^8$ | $1.6127 \times 10^8$ | $1.6835 \times 10^8$ |
| 1280 | $1.4514 \times 10^8$ | $1.6545 \times 10^8$ | $1.4875 \times 10^8$ | $1.6323 \times 10^8$ |
| 2560 | $1.5684 \times 10^8$ | $1.6532 \times 10^8$ | $8.3104 \times 10^7$ | $1.256 \times 10^8$ |
| 5120 | $6.957 \times 10^7$ | $1.5229 \times 10^8$ | $6.4447 \times 10^7$ | $8.7025 \times 10^7$ |
| 10240 | $1.366 \times 10^8$ | $6.9823 \times 10^7$ | $4.4714 \times 10^7$ | $4.054 \times 10^7$ |

##### *2.1.2. Neutralized viral titre estimated from viral titre with the most diluted antisera as viral titre*

Below, we estimate neutralized viral titre and use focus number in the eighth row as total viral titre. The neutralized viral titre in each well (Table S7) is obtained by substituting neutralized focus number in each well (Table S6) into the formula, $Viral\ titre = FN \times 20 \times 10^{3.5}\ FFU/ml$, where viral dilution factor is $10^{-3.5}$ and 50ul diluted virus in each well, $FN$ represents focus number in each well with one focus formation unit (FFU) representing one infectious influenza virus. Similarly, we provide total viral titre for each reference virus in Table S8.

**Table S6**. Neutralized foci number in each well (viral titre without the most diluted antisera as total viral titre)

| Serum Dilution | A/Victoria/653/2017 (cell grown) | A/Victoria/653/2017 (egg grown) | A/Switzerland /8060/2017(cell grown) | A/Switzerland /8060/2017( egg grown) |
|---|---|---|---|---|
| 80 | 1516 | 1928 | 2829 | 1928 |
| 160 | 1553 | 1963 | 2827 | 2017 |
| 320 | 1558 | 1960 | 2760 | 2029 |

| | | | | |
|---|---|---|---|---|
| 640 | 1545 | 1843 | 2516 | 2021 |
| 1280 | 1512 | 1645 | 1739 | 1940 |
| 2560 | 1510 | 607 | 1055 | 1345 |
| 5120 | 1304 | 312 | 214 | 735 |

**Table S7**. Neutralized viral titre in each well (FFU/ml, viral titre with the most diluted antisera as total viral titre)

| Serum Dilution | A/Victoria/653/2017 (cell grown) | A/Victoria/653/2017 (egg grown) | A/Switzerland /8060/2017(cell grown) | A/Switzerland /8060/2017( egg grown) |
|---|---|---|---|---|
| 80 | $9.588 \times 10^7$ | $1.2194 \times 10^8$ | $1.7892 \times 10^8$ | $1.2194 \times 10^8$ |
| 160 | $9.822 \times 10^7$ | $1.2415 \times 10^8$ | $1.7880 \times 10^8$ | $1.2757 \times 10^8$ |
| 320 | $9.8537 \times 10^7$ | $1.2396 \times 10^8$ | $1.7456 \times 10^8$ | $1.2833 \times 10^8$ |
| 640 | $9.7714 \times 10^7$ | $1.1656 \times 10^8$ | $1.5913 \times 10^8$ | $1.2781 \times 10^8$ |
| 1280 | $9.5627 \times 10^7$ | $1.0403 \times 10^8$ | $1.0998 \times 10^8$ | $1.2270 \times 10^8$ |
| 2560 | $9.5501 \times 10^7$ | $3.8390 \times 10^7$ | $6.6724 \times 10^7$ | $8.5065 \times 10^7$ |
| 5120 | $8.2472 \times 10^7$ | $1.9733 \times 10^7$ | $1.3535 \times 10^7$ | $4.6485 \times 10^7$ |

**Table S8**. Total viral titre for each reference virus (FFU/ml)

| A/Victoria/653/2017 (cell grown) | A/Victoria/653/2017 (egg grown) | A/Switzerland /8060/2017(cell grown) | A/Switzerland /8060/2017( egg grown) |
|---|---|---|---|
| $9.9549 \times 10^7$ | $1.2466 \times 10^8$ | $1.7987 \times 10^8$ | $1.2883 \times 10^8$ |

The median tissue culture infectious dose (TCID50), measuring viral concentration that infects 50% of susceptible cells, is a commonly used viral quantification assay to measure infectious influenza virus as used to measure viral titres in SC and MC assays, whereas, focus formation assay (FFA) is a rapid, reliable and reproducible viral quantification to infectious influenza virus (2). Since results obtained by TCID50 are linear to those obtained by FFA on $log_{10}$ scale (2), we assume $Y = aX + b$, where $Y$ represents viral titre obtained by TCID50 and $X$ represents viral titre obtained by FFA. By using FIT code in MATLAB 8.0 (The MathWorks, Inc., Natick, MA, US), we found $a = 1.182(95\%\ CI; 0.7857, 1.577)$ and $b = -2.854(-5.917, 0.2091)$ support the mutual conversion of results obtained by FFA and TCID50. Using $Y = aX + b$, we convert neutralized viral titre in FFU scale to TCID50 scale (Table S9 for cell control, Table S10 and Table S11 for viral titre with the most diluted antisera as total viral titre). Then, antibody concentrations at each serum dilution (80 to 10240 for cell control as total viral titre, 80 to 5120 for viral titre with the most diluted antisera as total viral titre) was estimated assuming 1mg/ml, providing antibodies concentrations for each dilution 50 ug/ml to 0.390625 ug/ml and for each dilution 50ug/ml to 0.78125ug/ml.

**Table S9**. Neutralized viral titre in each well (TCID50/ml, cell control as total viral titre)

| Serum Dilution | A/Sydney/22/2018 (cell grown) | A/Victoria/653/2017 (cell grown) | A/Victoria/653/2017 (egg grown) | A/Switzerland/8060/2017 (egg grown) |
|---|---|---|---|---|
| 80 | $1.1885 \times 10^{12}$ | $1.1812 \times 10^{12}$ | $1.1935 \times 10^{12}$ | $1.1397 \times 10^{12}$ |
| 160 | $1.1738 \times 10^{12}$ | $1.2116 \times 10^{12}$ | $1.2224 \times 10^{12}$ | $1.2124 \times 10^{12}$ |
| 320 | $1.2158 \times 10^{12}$ | $1.2158 \times 10^{12}$ | $1.2199 \times 10^{12}$ | $1.2224 \times 10^{12}$ |
| 640 | $1.2149 \times 10^{12}$ | $1.2050 \times 10^{12}$ | $1.1245 \times 10^{12}$ | $1.2158 \times 10^{12}$ |
| 1280 | $9.286 \times 10^{11}$ | $1.1779 \times 10^{12}$ | $9.710 \times 10^{11}$ | $1.1494 \times 10^{12}$ |
| 2560 | $1.63 \times 10^{11}$ | $1.1762 \times 10^{12}$ | $3.373 \times 10^{11}$ | $7.142 \times 10^{11}$ |
| 5120 | $3.7 \times 10^{10}$ | $1.0133 \times 10^{12}$ | $2.126 \times 10^{11}$ | $3.668 \times 10^{11}$ |
| 10240 | $1.27 \times 10^{11}$ | $2.459 \times 10^{11}$ | $1.095 \times 10^{11}$ | $9.16 \times 10^{10}$ |

**Table S10**. Neutralized viral titre in each well (TCID50/ml, viral titre with the most diluted antisera as total viral titre)

| Serum Dilution | A/Victoria/653/2017 (cell grown) | A/Victoria/653/2017 (egg grown) | A/Switzerland /8060/2017(cell grown) | A/Switzerland /8060/2017( egg grown) |
|---|---|---|---|---|
| 80 | $4.3734 \times 10^{11}$ | $6.7675 \times 10^{11}$ | $1.3578 \times 10^{12}$ | $6.7675 \times 10^{11}$ |
| 160 | $4.5692 \times 10^{11}$ | $6.9922 \times 10^{11}$ | $1.3561 \times 10^{12}$ | $7.3454 \times 10^{11}$ |
| 320 | $4.5959 \times 10^{11}$ | $6.9728 \times 10^{11}$ | $1.2983 \times 10^{12}$ | $7.4250 \times 10^{11}$ |
| 640 | $4.5265 \times 10^{11}$ | $6.2354 \times 10^{11}$ | $1.0974 \times 10^{12}$ | $7.3719 \times 10^{11}$ |
| 1280 | $4.3525 \times 10^{11}$ | $5.0726 \times 10^{11}$ | $5.6112 \times 10^{11}$ | $6.8441 \times 10^{11}$ |
| 2560 | $4.3420 \times 10^{11}$ | $8.2974 \times 10^{10}$ | $2.2641 \times 10^{11}$ | $3.5191 \times 10^{11}$ |
| 5120 | $3.3267 \times 10^{11}$ | $2.4777 \times 10^{10}$ | $1.2494 \times 10^{10}$ | $1.1745 \times 10^{11}$ |

**Table S11**. Total viral titre for each reference virus (TCID50/ml, viral titre with the most diluted antisera as total viral titre)

| A/Victoria/653/2017 (cell grown) | A/Victoria/653/2017 (egg grown) | A/Switzerland /8060/2017(cell grown) | A/Switzerland /8060/2017( egg grown) |
|---|---|---|---|
| $4.682 \times 10^{11}$ | $7.044 \times 10^{11}$ | $1.3709 \times 10^{12}$ | $7.478 \times 10^{11}$ |

#### 2.2. Estimation of virus neutralization parameters

We proposed three models to establish the quantitative relationship between neutralized viral titre and diluted antibody. They describe viral kinetics in a focus reduction assay; two with saturation and one without saturation. System 1 in main text describes saturation effect in both viral neutralization and antibody consumption, which is known as saturated neutralization, whereas System 4 describes saturation effect only in viral neutralization, which is known as semi-saturated neutralization.

$\begin{cases} \frac{dV(t)}{dt} = \frac{-\alpha A(t)V(t)}{1+\eta A(t)+\gamma V(t)} \\ \frac{dA(t)}{dt} = -\varphi A(t)V(t) \end{cases}$, (4), where $V(t)$ and $A(t)$ represents viral titre and antibody concentration with respect to time $t$. Parameters $\alpha$ represents virus neutralization rate by binding of antibodies; $\varphi$ represents antibody consumption rate by binding to virion; $\eta$ controls the saturation in the neutralization rate as antibody concentration increase; $\gamma$ controls the saturation in the neutralization rate as viral titre increase. Moreover, we propose System 2 in main text to describe unsaturated virus neutralization in one-hour incubation.

We use least-squares approach to obtain virus neutralization parameters. We have 8 datapoints $\{(A_1, V_1), \cdots, (A_8, V_8)\}$ for **cell control as total viral titre** and 7 datapoints $\{(A_1, V_1), \cdots, (A_7, V_7)\}$ for **virus titre with the most diluted antisera as total viral titre**, where $A_i$ represents diluted antibody concentration and $V_i$ represents neutralized virus titre. We defined the sum-of-squares error (SSE) as $SSE = \sum_{i=1}^{j} \left(log10(V_i) - log10(F(A_i))\right)^2$, where $V_i$ and $F(A_i)$ represent experimental and theoretical viral titre with respect to the i[th] diluted antibody concentration. $j = 8$ for cell control as total viral titre, and $j = 7$ for viral titre with the most diluted antisera as total viral titre. $F(A_i)$ is the integral of $\frac{dV(t)}{dt} = \frac{-\alpha A(t)V(t)}{1+\eta A(t)+\gamma V(t)}$ (saturated neutralization) or $\frac{dV(t)}{dt} = -\alpha A(t)V(t)$ (unsaturated neutralization) from 0 to 1, because influenza virus and diluted antibody are incubated for one hour.

We generated simulated data with different magnitudes of noise following Gaussian distribution $N(0,1)$. We fit System 1 and System 2 in main text to the simulated data. We found that Sum of Square Error (SSE) provided by System 1 in main text is smaller than that provided by System 2 in main text; namely, saturated virus neutralization is more accurate than unsaturated virus neutralization. Next, we also found that saturated virus neutralization is more robust to noise than unsaturated virus neutralization (data not shown).

##### *2.2.1. Estimation of virus neutralization parameters – FFU scale (Cell control as total virus titre)*

To estimate virus neutralization parameter (FFU scale) by fitting two proposed model with saturation to FRA data. We fit System 1 (main text) and System 4 to neutralized viral titre (FFU scale) and obtain virus neutralization parameters (Table S12 and S14, respectively). Because magnitude of parameters $\eta$ and $\varphi$ is very small, we use $\log_{10}$ transformation to fit both systems (Table S13 and S15, respectively). Because magnitude and category of virus neutralization parameter remain consistent between parameter provided System 1 (main text) or System 4, we use only the $\log_{10}$ transformation and System 1 (main text) in the rest text.

The estimated virus neutralization parameters from fitting System 2 (main text) to neutralized viral titre (FFU scale) are shown in Table S16. By comparing Table S12 to S16, SSE obtained from models with saturation effect is always 2/3 time smaller than SSE obtained from model without saturation effect, indicating a better bit of models with saturation.

**Table S12**. Estimated virus neutralization parameters (FFU scale, saturated neutralization)

| Dataset | $\alpha$ (ml/FFU/hour) | $\eta$ (ml/ug) | $\gamma$ (ml/FFU) | $\varphi$ (ml/ug/hour) | SSE | Category |
|---|---|---|---|---|---|---|
| A/Sydney/22/2018 (cell grown) | 8.0161 | $7.268 \times 10^{-7}$ | 2.1177 | $1.005 \times 10^{-6}$ | 1.0459 | Small |
| A/Victoria/653/2017 (cell grown) | 38.7690 | 0.2458 | 5.1029 | 1.2023 | 0.0072 | Large |
| A/Victoria/653/2017 (egg grown) | 13.9600 | $2.325 \times 10^{-6}$ | 2.8163 | $3.215 \times 10^{-6}$ | 0.2516 | Small |
| A/Switzerland/8060/2017 (egg grown) | 6.8092 | 0.0053 | 0.0588 | 0.0086 | 0.2721 | Large |

**Table S13**. Estimated virus neutralization parameters (FFU scale, saturated neutralization, log10 parameter estimation)

| Dataset | $\alpha$ (ml/FFU/hour) | $\eta$ (ml/ug) | $\gamma$ (ml/FFU) | $\varphi$ (ml/ug/hour) | SSE | Category |
|---|---|---|---|---|---|---|
| A/Sydney/22/2018 (cell grown) | 8.0161 | $1.4537 \times 10^{-7}$ | 2.1177 | $2.0112 \times 10^{-7}$ | 1.0459 | Small |
| A/Victoria/653/2017 (cell grown) | 8.8695 | 0.0097 | 0.1849 | 0.0177 | 0.0287 | Large |
| A/Victoria/653/2017 (egg grown) | 13.69 | $2.3258 \times 10^{-6}$ | 2.8164 | $3.214 \times 10^{-6}$ | 0.2516 | Small |
| A/Switzerland/8060/2017 (egg grown) | 11.5518 | $2.0881 \times 10^{-6}$ | 1.9678 | $3.031 \times 10^{-6}$ | 0.0907 | Small |

**Table S14**. Estimated virus neutralization parameters (FFU scale, semi-saturated neutralization)

| Dataset | $\alpha$ (ml/FFU/hour) | $\eta$ (ml/ug) | $\gamma$ (ml/FFU) | $\varphi$ (ml/ug/hour) | SSE | Category |
|---|---|---|---|---|---|---|
| A/Sydney/22/2018 (cell grown) | 8.0162 | $9.0862 \times 10^{-7}$ | 2.1177 | $1.981 \times 10^{-6}$ | 1.0459 | Small |
| A/Victoria/653/2017 (cell grown) | 12.8914 | 0.0293 | 0.0038 | 0.2584 | 0.0445 | Large |
| A/Victoria/653/2017 (egg grown) | 13.6901 | $2.3258 \times 10^{-6}$ | 2.8163 | $4.493 \times 10^{-6}$ | 0.2516 | Small |
| A/Switzerland/8060/2017 (egg grown) | 11.5519 | $2.0880 \times 10^{-6}$ | 1.9678 | $4.027 \times 10^{-6}$ | 0.0907 | Small |

**Table S15**. Estimated virus neutralization parameters (FFU scale, semi-saturated neutralization, log10 parameter estimation)

| Dataset | $\alpha$ (ml/FFU/hour) | $\eta$ (ml/ug) | $\gamma$ (ml/FFU) | $\varphi$ (ml/ug/hour) | SSE | Category |
|---|---|---|---|---|---|---|
| A/Sydney/22/2018 (cell grown) | 8.0161 | $1.4537 \times 10^{-7}$ | 2.1177 | $2.0112 \times 10^{-7}$ | 1.0459 | Small |
| A/Victoria/653/2017 (cell grown) | 8.8695 | 0.0097 | 0.1849 | 0.0177 | 0.0287 | Large |
| A/Victoria/653/2017 (egg grown) | 13.69 | $2.3258 \times 10^{-6}$ | 2.8164 | $3.2154 \times 10^{-6}$ | 0.2516 | Small |
| A/Switzerland/8060/2017 (egg grown) | 11.5518 | $2.0881 \times 10^{-6}$ | 1.9678 | $3.0314 \times 10^{-6}$ | 0.0907 | Small |

**Table S16**. Estimated virus neutralization parameters (FFU scale, unsaturated neutralization)

| Dataset | $\alpha$ (ml/FFU/hour) | $\varphi$ (ml/ug/hour) | SSE | Category |
|---|---|---|---|---|
| A/Sydney/22/2018 (cell grown) | 4.3064 | $5.3983 \times 10^{-7}$ | 1.5675 | Small |
| A/Victoria/653/2017 (cell grown) | 7.9506 | 0.0091 | 0.0330 | Large |
| A/Victoria/653/2017 (egg grown) | 6.5487 | $2.5847 \times 10^{-6}$ | 0.4896 | Small |

| | | | | |
|---|---|---|---|---|
| A/Switzerland/8060/2017 (egg grown) | 6.5497 | 0.0066 | 0.2744 | Large |

*2.2.2. Estimation of virus neutralization parameters – TCID50 scale (Cell control as total virus titre)*

We fit System 1 (main text) and System 2(main text) to neutralized viral titre (TCID50 scale) and obtain virus neutralization parameters (Table S16 and S17, respectively). By comparing Table S17 to S18, SSE obtained from models with saturation effect is always 2/3 time smaller than SSE obtained from model without saturation effect, indicating a better bit of models with saturation.

**Table S17.** Estimated virus neutralization parameters (TCID50 scale, saturated neutralization, log10 parameter estimation)

| Data | $\alpha$ (ml/TCID50/hour) | $\eta$ (ml/ug) | $\gamma$ (ml/TCID50) | $\varphi$ (ml/ug/hour) | $SSE^{*}$ | Category |
|---|---|---|---|---|---|---|
| A/Sydney/22/2018 (cell grown) | 6.4228 | $2.376 \times 10^{-7}$ | 1.7591 | $3.312 \times 10^{-7}$ | 1.8604 | Small |
| A/Victoria/653/2017 (cell grown) | 8.1600 | 0.0112 | $3.251 \times 10^{-4}$ | 0.0265 | 0.0602 | Large |
| A/Victoria/653/2017 (egg grown) | 11.8852 | $6.253 \times 10^{-7}$ | 2.5368 | $8.719 \times 10^{-7}$ | 0.4482 | Small |
| A/Switzerland/8060/2017 (egg grown) | 10.1114 | $5.736 \times 10^{-7}$ | 1.7609 | $8.454 \times 10^{-7}$ | 0.1550 | Small |

**Table S18**. Estimated virus neutralization parameters (TCID50 scale, unsaturated neutralization)

| Dataset | $\alpha$ (ml/TCID50/hour) | $\varphi$ (ml/ug/hour) | SSE | Category |
|---|---|---|---|---|
| A/Sydney/22/2018 (cell grown) | 3.5890 | $8.5461 \times 10^{-7}$ | 2.7459 | Small |
| A/Victoria/653/2017 (cell grown) | 7.4322 | 0.0072 | 0.0528 | Large |
| A/Victoria/653/2017 (egg grown) | 5.9588 | $8.4239 \times 10^{-5}$ | 0.8726 | Small |
| A/Switzerland/8060/2017 (egg grown) | 6.5716 | 0.0350 | 0.4938 | Large |

##### *2.2.3. Neutralization parameters of A/H3N2 viruses- FFU scale (cell control as total viral titre)*

Virus neutralization parameters were estimated for FRA data obtained for thirteen A/H3N2 viruses selected from 49 virus circulating during 2014–2019 as described in Section 2.1.1. The only standard to select virus is the focus number in well without antisera is higher than that with antisera. The tested virus used in FRA is MDCK-SIAT2 cell-grown A/Tasmania/512/2019, MDCK-SIAT1 cell-grown A/Christchurch/514/2019, MDCK-SIAT1 cell-grown A/Christchurch/518/2019, MDCK-SIAT1 cell-grown A/South Australia/301/2019, MDCK-SIAT2 cell-grown A/Victoria/2159/2019, MDCK-SIAT2 cell-grown A/Sydney/770/2019, MDCK-SIAT2 cell-grown A/Sydney/500/2019, MDCK-SIAT1 cell-grown A/Canberra/108/2019, MDCK-SIAT1 cell-grown A/Fiji/25/2019, MDCK-SIAT1 cell-grown A/Fiji/36/2019, Egg-grown A/Victoria/653/2017, and Egg-grown A/Switzerland/8060/2017 (Ab ON). Selected viruses are listed in Fig. S5. Similar to A/Brisbane/32/2017, we found that the estimated virus neutralization parameters can classified into two categories, large and small antibody consumption, however the majority were small. The magnitude of virus neutralization parameter remained consistent among the thirteen influenza virus (data not shown).

We found that Pearson correlation coefficient between $log10(\eta)$ and $log10(\varphi)$ is 0.9842, indicating that antibody saturation and antibody consumption are strongly positively correlated (Fig. S6); this warrants further study. This remains consistent with results provided by virus titre with the most diluted antisera. One possible explanation is that because the objective function may not be convex, there may be many local

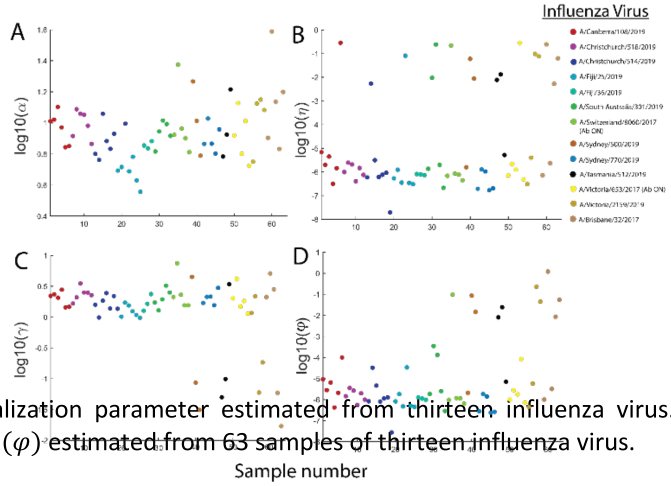


**Figure S5**. Four virus neutralization parameter estimated from thirteen influenza virus. A-D) represent $log_{10}(\alpha)$, $log_{10}(\eta)$, $log_{10}(\gamma)$ and $log_{10}(\varphi)$ estimated from 63 samples of thirteen influenza virus.

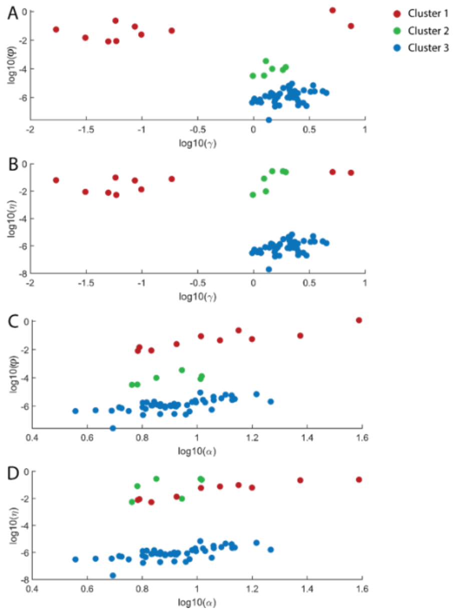


minima and if we start from different points the minimisation algorithm can converge to various local minima.

##### *2.2.4. Estimation of virus neutralization parameters – FFA scale (most diluted antisera as total virus titre)*

We fit System 1 (main text) and System 2 (main text) to neutralized viral titre (FFU scale) and obtain virus neutralization parameters (Table S19 and Table S20, respectively). Similarly, the virus neutralization parameter is divided into two categories, including large and small antibody consumption rate. By comparing Table S19 to S20, SSE obtained from models with saturation effect is always 2/3 time smaller than SSE obtained from model without saturation effect, indicating a better bit of models with saturation.

**Figure S6**. Antibody saturation and antibody consumption are strongly positively correlated. A) represents $log_{10}(\gamma)$ plotted against $log_{10}(\varphi)$, B) represents $log_{10}(\gamma)$ plotted against $log_{10}(\eta)$, C) represents $log_{10}(\alpha)$ plotted against $log_{10}(\varphi)$ and D) represents $log_{10}(\alpha)$ plotted against $log_{10}(\eta)$.

**Table S19.** Estimated virus neutralization parameters (FFU scale, saturated virus neutralization)

| Dataset | $\alpha$ (ml/FFU/hour) | $\eta$ (ml/ug) | $\gamma$ (ml/FFU) | $\varphi$ (ml/ug/hour) | $SSE^{*}$ | Category |
|---|---|---|---|---|---|---|

| | | | | | | |
|---|---|---|---|---|---|---|
| A/Victoria/653/2017 (cell grown) | 96.8327 | 0.7788 | 14.2576 | 7.167 | 0.0111 | Large |
| A/Victoria/653/2017 (egg grown) | 24.6437 | 0.9199 | 5.2143 | 0.5620 | 0.2854 | Large |
| A/Switzerland/8060/2017(cell grown) | 3.8399 | $1.5 \times 10^{-6}$ | 0.6529 | $2.16 \times 10^{-7}$ | 0.077 | Small |
| A/Switzerland/8060/2017(cell grown) | 6.3978 | $2.44 \times 10^{-7}$ | 0.9358 | $3.63 \times 10^{-7}$ | 0.0579 | Small |

**Table S20.** Estimated virus neutralization parameters (FFU scale, unsaturated neutralization)

| Dataset | $\alpha$ (ml/TCID50/hour) | $\varphi$ (ml/ug/hour) | SSE | Category |
|---|---|---|---|---|
| A/Victoria/653/2017 (cell grown) | 57.8936 | 5.4794 | 0.0253 | Large |
| A/Victoria/653/2017 (egg grown) | 2.9129 | 0.0138 | 0.4356 | Large |
| A/Switzerland/8060/2017(cell grown) | 2.4678 | $1.9077 \times 10^{-7}$ | 0.3444 | Small |
| A/Switzerland/8060/2017(cell grown) | 8.1858 | 0.5757 | 0.1808 | Large |

##### *2.2.5. Estimation of virus neutralization parameters – TCID50 scale (most diluted antisera as total virus titre)*

We fit System 1 (main text) and System 2 (main text) to neutralized viral titre (TCID50 scale) and obtain virus neutralization parameters (Table S21 and Table S22, respectively). Similarly, the virus neutralization parameter is divided into two categories, including large and small antibody consumption rate. By comparing Table S21 to S22, SSE obtained from models with saturation effect is always 2/3 time smaller than SSE obtained from model without saturation effect, indicating a better bit of models with saturation.

**Table S21.** Estimated virus neutralization parameters (TCID50 scale)

| Dataset | $\alpha$ (ml/TCID50/hour) | $\eta$ (ml/ug) | $\gamma$ (ml/TCID50) | $\varphi$ (ml/ug/hour) | $SSE^{*}$ | Category |
|---|---|---|---|---|---|---|
| A/Victoria/653/2017 (cell grown) | 79.0081 | 3.6175 | 1.6091 | 1.7164 | 0.0520 | Large |
| A/Victoria/653/2017 (egg grown) | 4.1281 | $5.1898 \times 10^{-7}$ | 0.7571 | $7.9082 \times 10^{-6}$ | 0.3745 | Small |
| A/Switzerland/8060/2017(cell grown) | 3.2653 | $6.9899 \times 10^{-7}$ | 0.5555 | $1.0572 \times 10^{-7}$ | 0.1270 | Small |

| | | | | | | |
|---|---|---|---|---|---|---|
| A/Switzerland/8060/2017(cell grown ) | 5.7636 | 0.1151 | $7.0757 \times 10^{-6}$ | 0.0757 | 0.3194 | Large |

**Table S22.** Estimated virus neutralization parameters (TCID50 scale, unsaturated neutralization)

| Dataset | $\alpha$ (ml/TCID50/hour) | $\varphi$ (ml/ug/hour) | SSE | Category |
|---|---|---|---|---|
| A/Victoria/653/2017 (cell grown) | 27.7330 | 1.6666 | 0.0484 | Large |
| A/Victoria/653/2017 (egg grown) | 8.2820 | 0.5307 | 0.9333 | Large |
| A/Switzerland/8060/2017(cell grown) | 65.9908 | 5.1020 | 0.8763 | Large |
| A/Switzerland/8060/2017(cell grown) | 7.7685 | 0.3882 | 0.3283 | Large |

##### *2.2.6. Neutralization parameters of A/H3N2 viruses- FFU scale (most diluted antisera as total viral titre)*

Four virus neutralization parameters were estimated for FRA data obtained 49 virus circulating during 2014–2019 as described in Section 2.1.1. The only standard to select virus is that the focus number in well from the first to seventh row is higher than that in eighth row. Similarly, we found that the estimated virus neutralization parameters can classified into two categories, large and small antibody consumption, however the majority were small. Moreover, by plotting amino acid sequence distance of HA between test and reference virus against antibody consumption, two categories by antibody consumption is independent of amino acid distance of HA between test and reference virus (shown in Fig. S7). The amino acid distance of HA between test and reference virus is computed on MEGA X(Molecular evolutionary genetics analysis across computing platforms) [6].

Next, we use Pearson correlation to analyse four parameter of virus neutralization kinetics, including virus neutralization, antibody consumption, antibody saturation and virus saturation (shown in Fig. S8). Our result suggests that a positive correlation between antibody saturation ($log10(\eta)$) and antibody consumption ($log10(\varphi)$) (Pearson correlation coefficient, 0.7601, Fig. S8A, B, D and D) and between virus neutralization ($log10(\alpha)$)and antibody consumption ($log10(\varphi)$) (correlation coefficient, 0.7770; S8F), showing that the more efficient the antibody is, the more antibody is consumed.

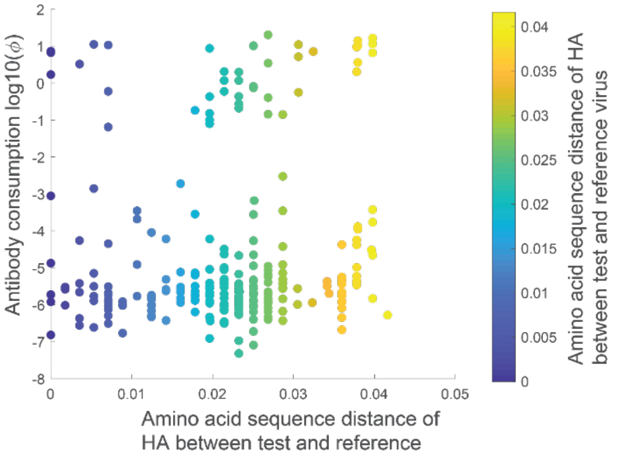


**Figure S7.** Two categories of antibody consumption are independent of amino acid distance (HA) between test and reference virus.

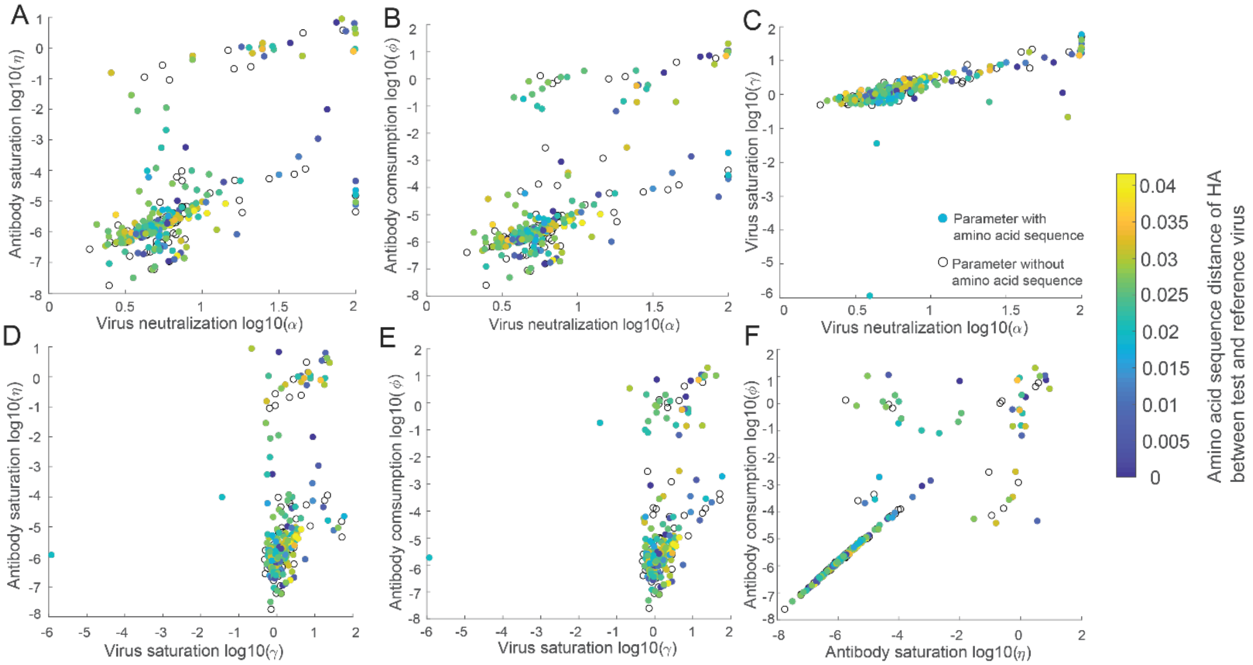


**Figure S8.** Comparison of four virus neutralization parameters, including virus neutralization, antibody consumption, antibody saturation and virus saturation. Antibody saturation and antibody consumption are strongly positively correlated, A) and B), D) and E). A) virus neutralization is plotted against antibody saturation. A) virus neutralization is plotted against antibody saturation. A) virus neutralization is plotted against antibody saturation; B) virus neutralization is plotted against antibody consumption; C) virus neutralization is plotted against virus saturation; D) virus saturation is plotted against antibody saturation; E) virus saturation is plotted against antibody consumption; F) antibody saturationis plotted against antibody consumption.

### 3. Bistable growth kinetics of influenza virus in the presence of neutralizing antibody

#### 3.1. Virus neutralization parameter estimated from viral titre with the most diluted antisera as total viral titre

##### *3.1.1. Antibody concentration kinetics with A/H7N9 viral replication parameter in presence of antibody (ODE models without eclipse phase)*

First, we substitute H7N9 virus replication (Table S1) and neutralization parameter obtained from dataset 3 with **small antibody consumption rate** (Table S21) into System 3 (main text). Due to small antibody consumption rate, the amount of consumed antibody is very small and then viral kinetics with decreasing antibody concentration is approximated by viral kinetics with constant antibody concentration (Fig. S9).

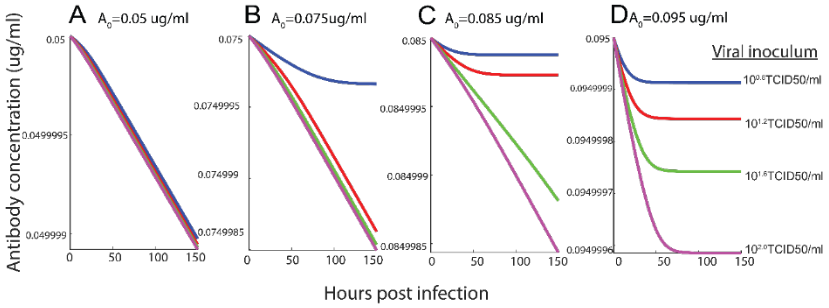


**Figure S9.** Antibody concentration kinetics with different inoculum sizes in 144-hour incubation. A-D) represents different initial antibody concentration 0.05ug/ml, 0.075ug/ml, 0.085ug/ml and 0.095ug/ml.

Then, we substitute H7N9 virus replication (Table S1) and neutralization parameter obtained from dataset 1 with **large antibody consumption rate** (Table S21) into System 3 (main text). Due to large antibody consumption rate, viral survival corresponds to depletion of antibody; viral eradication coincides with antibody remaining, shown in Fig. S10 and Fig. 3 (main text).

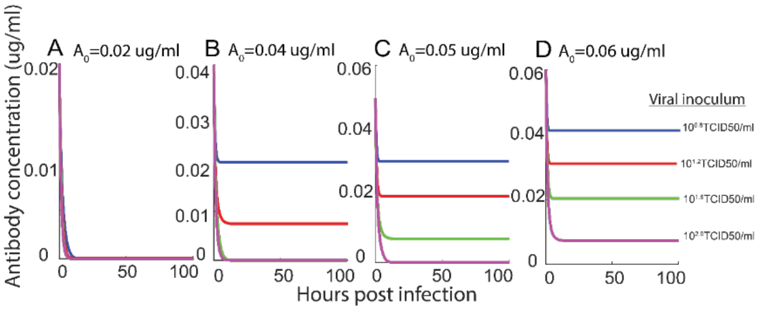


**Figure S10.** Antibody concentration kinetics with different inoculum sizes in 100-hour incubation. A-D) represents different initial antibody concentration 0.02ug/ml, 0.04ug/ml, 0.05ug/ml and 0.06ug/ml.

##### *3.1.2. Bistable kinetics of A/H7N9 with small antibody consumption and eclipse phase (DDE model)*

We introduce the eclipse phase to a two-dimensional deterministic model for the rate of change of the viral titre $V(TCID\,50/ml)$ and describe how this rate is determined by antibody concentration $A(ug/ml)$,

$$\begin{cases} \frac{dV(t)}{dt} = \frac{\rho V(t-\tau)}{1+\beta V(t)} - \sigma V(t) - \frac{\alpha A(t)V(t)}{1+\eta A(t)+\gamma V(t)} \\ \frac{dA(t)}{dt} = \frac{-\varphi A(t)V(t)}{1+\eta A(t)+\gamma V(t)} \end{cases}, (5).$$

In this section, we integrated H7N9 virus replication parameter (Table S1) with eclipse phase and virus neutralization parameter obtained from dataset 3 with **small antibody consumption rate** (Table S21) into System 5.

Antibody-induced bistable viral kinetics exists. Due to small antibody consumption rate, antibody concentration is approximated as constant during 144 hours incubation (Fig. S11), and then initial antibody concentration is used as constant antibody concentration. Specifically, for antibody concentration $A = 0.092\,ug/ml$, virus with inoculum $10^{1.2}TCID\,50/ml$, $10^{1.6}TCID\,50/ml$ and $10^{2.0}TCID\,50/ml$ survives (Fig. S12B and Fig. S12E); for antibody concentration $A = 0.1\,ug/ml$, virus with inoculum $10^{1.6}TCID\,50/ml$ and $10^{2.0}TCID\,50/ml$ survives (Fig. S12C and Fig. S12E). For antibody concentration less than $A_1$, virus survives independent of viral inocula (Fig. S12A and Fig. S12E). For antibody concentration higher than $A_2$, virus is inhibited independent of viral inoculum (Fig. S12D and Fig. S12E).

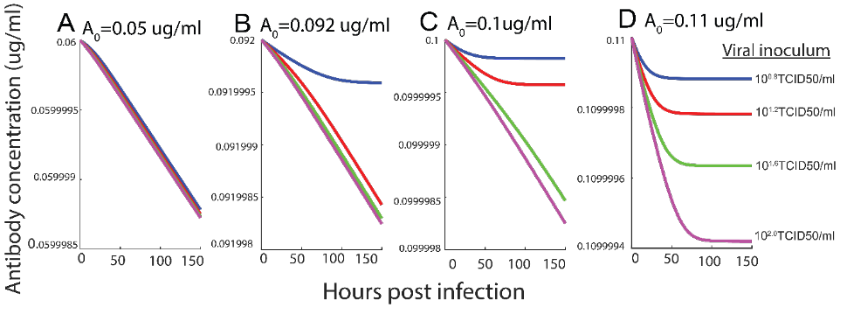


**Figure S11**. Antibody concentration kinetics with small antibody consumption rate. A-D) represents antibody concentration kinetics with initial antibody concentration $0.05\,ug/ml$, $0.092\,ug/ml$, $0.1\,ug/ml$ and $0.11\,ug/ml$.

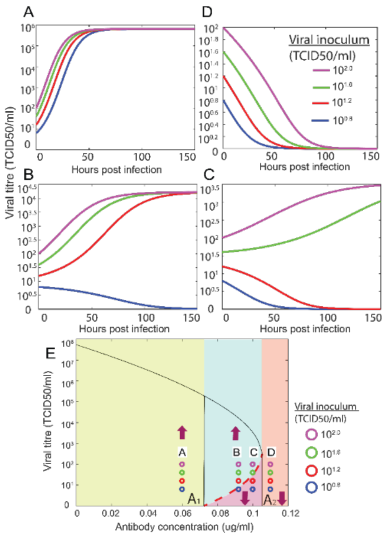


**Figure S12**. Simulated kinetics of H7N9 virus (with eclipse phase) with different combinations of inoculum sizes and antibody concentrations with small antibody consumption rate (dataset 3). Bifurcation diagram E) showing viral titre as a function of antibody concentration. Viral inoculum threshold increases with increase of antibody concentration (red dashed curve). The maximal capacity of viral titre decreases with increases of antibody concentration (black solid curve). Purple arrows represent any viral inoculum size. A) When antibody concentration is between 0 and $A_1$ virus with any viral inoculum survive. D) Virus with any inoculum size is inhibited when antibody concentration is greater than $A_2$. B) and C) When antibody concentration is between $A_1$ and $A_2$ virus survives if viral inoculum is above dashed red curve and is inhibited if viral inoculum is below dashed red curve. Purple, green, red, blue curve and circle represents viral kinetics with viral inoculum $10^{0.8}$, $10^{1.2}$, $10^{1.6}$ and $10^{2.0}$TCID50/ml in (A-E).

##### *3.1.3. Bistable kinetics of A/H7N9 with large antibody consumption and eclipse phase (DDE model)*

Then, we substitute H7N9 virus replication (Table S2) and neutralization parameter obtained from dataset 3 with **large antibody consumption rate** (Table S21) into System 5. Antibody-induced bistable viral kinetics exists.

With initial antibody concentration $A_0 = 0.03\,ug/ml$, virus survives independent on viral inoculum and antibody is depleted (Fig. S13A and Fig. S14A). With initial antibody concentration $A_0 = 0.04\,ug/ml$, virus with viral inoculum $10^{1.2} TCID\,50/ml$, $10^{1.6} TCID\,50/ml$ and $10^{2.0} TCID\,50/ml$ converges to maximum capacity; meanwhile corresponding antibody concentration is depleted (Fig. S13B and Fig.S14B). With initial antibody concentration $A_0 = 0.05\,ug/ml$, only virus with viral inoculum $10^{1.6} TCID\,50/ml$ and $10^{2.0} TCID\,50/ml$ survive and converge to maximum capacity (Fig. S13C and Fig.S14C). With initial antibody concentration $A_0 = 0.07\,ug/ml$, virus is inhibited independent of viral inoculum and antibody remains (Fig. S13D and Fig.S14D).

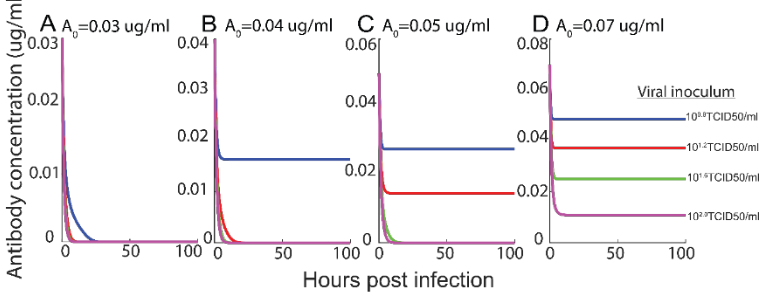


**Figure S13**. Antibody concentration kinetics with different inoculum sizes in 100-hour incubation**.** A-D) represents antibody concentration kinetics with different initial antibody concentration $0.03\,ug/ml$, $0.04\,ug/ml$, $0.05\,ug/ml$ and $0.07\,ug/ml$.

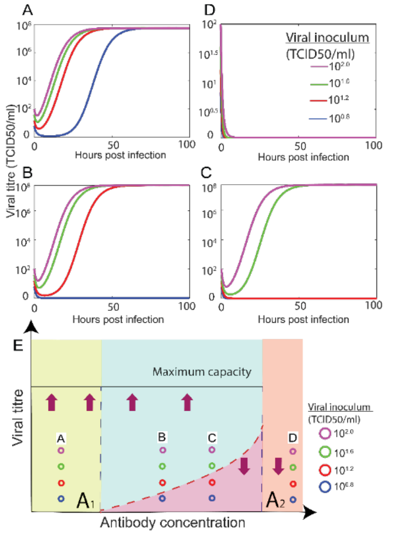


**Figure S14.** Simulated kinetics of H7N9 virus (with eclipse phase) with different combinations of inoculum sizes and antibody concentrations with large antibody consumption rate (dataset 1). Viral survival corresponds to antibody depletion and viral eradication coincides with antibody existence. Bifurcation diagram E) showing viral titre as a function of initial antibody concentration (schematic diagram). Viral inoculum threshold increases with increase of antibody concentration (dashed red line). Maximum capacity of viral titre remains constant if virus survives (black solid line). Purple arrows represent any viral inoculum size. A) When antibody concentration is between 0 and $A_1$, virus with any viral inoculum survive. B) and C) When antibody concentration is between $A_1$ and $A_2$ , viral kinetics survives if viral inoculum is above dashed red curve and is inhibited if viral inoculum is below dashed red curve. D) Virus with any inoculum size is inhibited when antibody concentration is greater than $A_2$. Purple, green, red, blue curve and circle represent viral kinetics with viral inoculum $10^{0.8}$, $10^{1.2}$, $10^{1.6}$ and $10^{2.0}$TCID50/ml in (A-E).

#### 3.2. Virus neutralization parameter estimated from cell control as total virus titre

##### *3.2.1. Bistable kinetics of A/H1N1pdm09 with small and large antibody consumption rate (ODE model without eclipse)*

In this section, we substitute H1N1pdm09 virus replication (Table S1) and neutralization parameter obtained from dataset 3 with **small antibody consumption rate** (Table S17) into System 3 (main text). Antibody-induced bistable viral kinetics exists.

Due to small antibody consumption rate, antibody concentration is approximated as constant during 144 hours incubation (Fig. S15), and then initial antibody concentration is used as constant antibody concentration. Then, at low antibody concentration $A = 0.03\, ug/ml$, virus with inoculum $10^{1.6} TCID\, 50/ml$ and $10^{2.0} TCID\, 50/ml$ survive (Fig. S16B and S16E), whereas at high antibody concentration $A = 0.035\, ug/ml$, only virus with high inoculum $10^{2.0} TCID\, 50/ml$ survives (Fig. S16C and S16E). At antibody concentration less than the threshold $A_1$, the virus survives independent of inoculum (Fig. S16A and S16E), and at antibody concentration higher than the threshold $A_2$, the virus is inhibited independent of inoculum size (Fig. S16D and S16E).

Then, we substitute H1N1pdm09 virus replication (Table S1) and neutralization parameter obtained from dataset 2 with **large antibody consumption rate** (Table S17) into System 3 (main text). Antibody-induced bistable viral kinetics exists. With initial antibody concentration $A_0 = 0.02\, ug/ml$, virus with inoculum $10^{1.2} TCID\, 50/ml$, $10^{1.6} TCID\, 50/ml$ and $10^{2.0} TCID\, 50/ml$ survives, while antibody is depleted (Fig. S18B, Fig. S17E and Fig. S17). With initial antibody concentration $A_0 = 0.025\, ug/ml$, only virus with inoculum $10^{1.6} TCID\, 50/ml$ and $10^{2.0} TCID\, 50/ml$ survives (Fig. S18C and Fig.S18E).

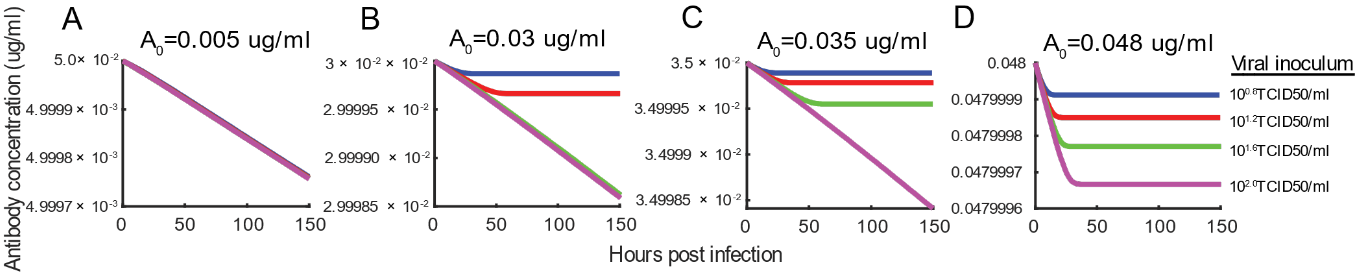


**Figure S15**. Antibody concentration kinetics with different inoculum sizes in 144-hour incubation. A-D) represents different initial antibody concentration 0.005ug/ml, 0.03ug/ml, 0.035ug/ml and 0.048ug/ml.

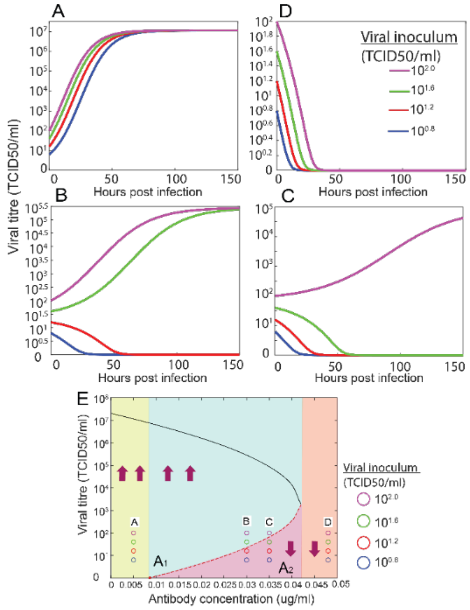


**Figure S16**. Simulated kinetics of H1N1pdm09 virus with different combinations of inoculum sizes and antibody concentrations with low antibody consumption (dataset 3). Bifurcation diagram (E) showing viral titre as a function of antibody concentration. Viral inoculum threshold increases with increase of antibody concentration (red dashed curve). The maximal capacity of viral titre decreases with increases of antibody concentration (black solid curve). Purple arrows represent any viral inoculum size. A) When antibody concentration is between 0 and $A_1$ virus with any viral inoculum survive. B) Virus with any inoculum size is inhibited when antibody concentration is greater than $A_2$. C) and D) when antibody concentration is between $A_1$ and $A_2$ virus survives if viral inoculum is above dashed red curve and is inhibited if viral inoculum is below dashed red curve. Purple, green, red and blue lines and circles represent viral kinetics with viral inoculum $10^{0.8}$, $10^{1.2}$, $10^{1.6}$ and $10^{2.0}$ TCID50/ml in A–D, also shown as virus inoculum in E.

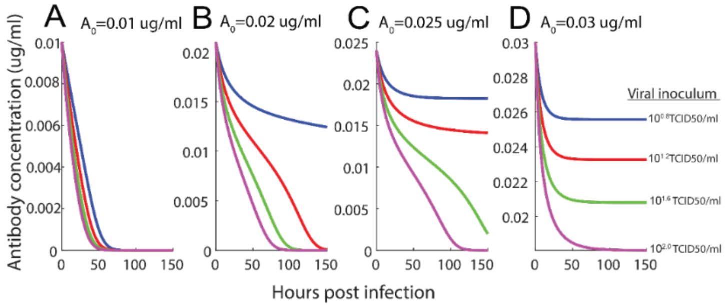


**Figure S17**. Antibody concentration kinetics with different inoculum sizes in 144-hour incubation. A-D) represents different initial antibody concentration 0.01ug/ml, 0.02ug/ml, 0.025ug/ml and 0.03ug/ml.

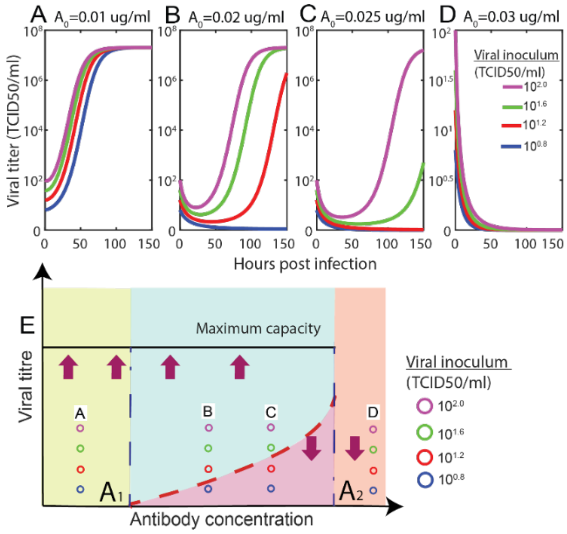


**Figure S18**. Simulated kinetics of H1N1pdm09 virus with different combinations of inoculum sizes and antibody concentrations with high antibody consumption (dataset 2). Viral survival corresponds to antibody depletion and viral eradication coincides with antibody existence. Bifurcation diagram (E) showing viral titre as a function of initial antibody concentration (schematic diagram). Viral inoculum threshold increases with increase of antibody concentration (dashed red line). Maximum capacity of viral titre remains constant if virus survives (black solid line). Purple arrows represent any viral inoculum size. A) When antibody concentration is between 0 and $A_1$, virus with any viral inoculum survive. B) and C) When antibody concentration is between $A_1$ and $A_2$ , viral kinetics survives if viral inoculum is above dashed red curve and is inhibited if viral inoculum is below dashed red curve. D) Virus with any inoculum size is inhibited when antibody concentration is greater than $A_2$. Purple, green, red and blue lines and circles represent viral kinetics with viral inoculum $10^{0.8}$, $10^{1.2}$, $10^{1.6}$ and $10^{2.0}$ TCID50/ml in A–D, also shown as virus inoculum in E.

##### *3.2.2. Bistable kinetics of A/H7N9 with small and large antibody consumption rate (ODE model without eclipse)*

We integrate H7N9 virus replication (Table S1) and neutralization parameters obtained from dataset 3 (**small antibody consumption rate**) (Table S17) into System 3 (main text) demonstrated the existence of antibody-induced bistable viral kinetics. Antibody-induced bistable viral kinetics exists. Due to small antibody consumption rate, antibody concentration is approximated as constant during 144 hours incubation (Fig. S19), and then initial antibody concentration is used as constant antibody concentration. Specifically, for antibody concentration $A = 0.05\, ug/ml$, virus with inoculum $10^{1.2} TCID\ 50/ml$, $10^{1.6} TCID\ 50/ml$and $10^{2.0} TCID\ 50/ml$ survives (Fig. S20B and Fig.S20E); for antibody concentration $A = 0.06\, ug/ml$, only virus with inoculum $10^{1.6} TCID\ 50/ml$ and $10^{2.0} TCID\ 50/ml$ survives (Fig. S20C and Fig.S20E). For antibody concentration less than $A_1$, virus survives independent of viral inoculum (Fig. S20A and Fig. S20E). For antibody concentration higher than $A_2$, virus is inhibited independent of viral inoculum (Fig. S20D and Fig. S20E).

By substituting H7N9 virus replication (Table S1) and neutralization parameter obtained from dataset 2 **large antibody consumption rate** (Table S17) into System 3 (main text), antibody-induced bistable viral kinetics also exists. With initial antibody concentration $A_0 = 0.01\,ug/ml$, virus survives independent on viral inoculum and antibody is depleted (Fig. S21A and Fig. S22A). With initial antibody concentration $A_0 = 0.034\,ug/ml$, virus with viral inoculum $10^{1.2}TCID\,50/ml$, $10^{1.6}TCID\,50/ml$ and $10^{2.0}TCID\,50/ml$ converges to maximum capacity; meanwhile corresponding antibody concentration is depleted (Fig. S21B and Fig.S22B). With initial antibody concentration $A_0 = 0.04\,ug/ml$, only virus with viral inoculum $10^{2.0}TCID\,50/ml$ survives and converges to maximum capacity (Fig. S21C and Fig.S22C). With initial antibody concentration $A_0 = 0.05\,ug/ml$, virus is inhibited independent of viral inoculum and antibody remains (Fig. S21D and Fig.S22D).

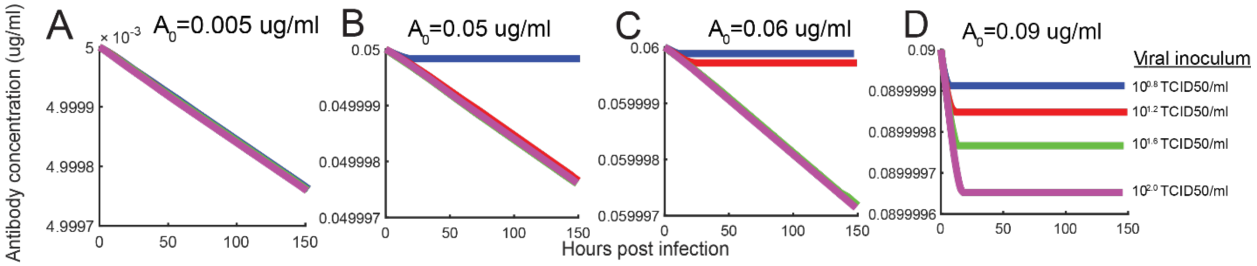


**Figure S19.** Antibody concentration kinetics with small antibody consumption rate. (A-D) represents different initial antibody concentration 0.005ug/ml, 0.05ug/ml, 0.06ug/ml and 0.09ug/ml**.**

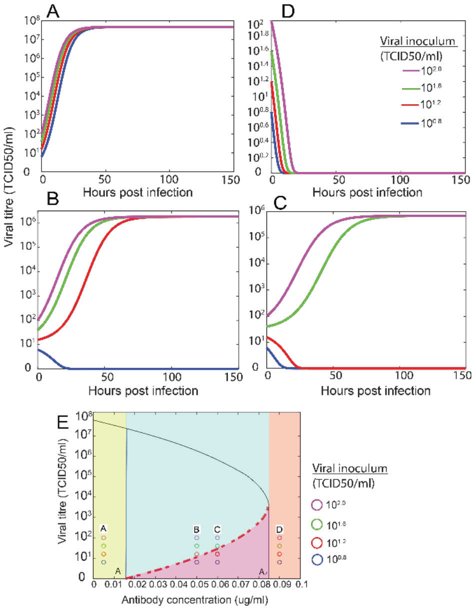


**Figure S20**. Simulated kinetics of H7N9 virus with different combinations of inoculum sizes and antibody concentrations with small antibody consumption rate (dataset 3). Bifurcation diagram E) showing viral titre as a function of antibody concentration. Viral inoculum threshold increases with increase of antibody concentration (red dashed curve). The maximal capacity of viral titre decreases with increases of antibody concentration (black solid curve). Purple arrows represent any viral inoculum size. A) When antibody concentration is between 0 and $A_1$ virus with any viral inoculum survive. D) Virus with any inoculum size is inhibited when antibody concentration is greater than $A_2$. B) and C) When antibody concentration is between $A_1$ and $A_2$ virus survives if viral inoculum is above dashed red curve and is inhibited if viral inoculum is below dashed red curve. Purple, green, red, blue curve and cycle represent viral kinetics with viral inoculum $10^{0.8}$, $10^{1.2}$, $10^{1.6}$ and $10^{2.0}$ TCID50/ml in (A-E).

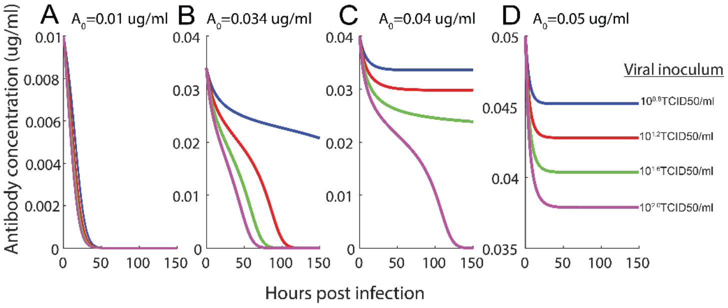


**Figure S21**. Antibody concentration kinetics with different inoculum sizes in 144-hour incubation**.** A-D) represents antibody concentration kinetics with different initial antibody concentration $0.01\,ug/ml$, $0.034\,ug/ml$, $0.04\,ug/ml$ and $0.05\,ug/ml$

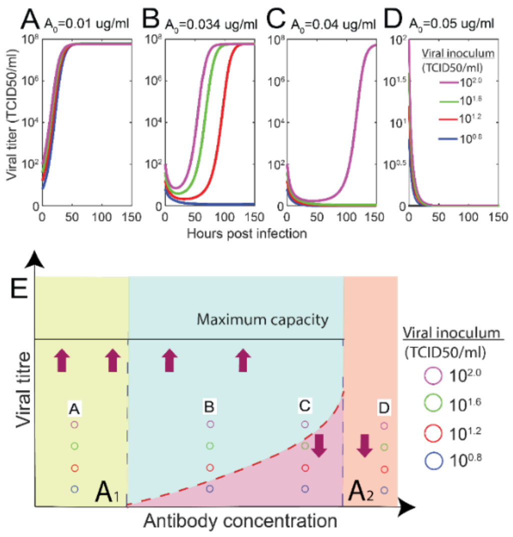


**Figure S22.** Simulated kinetics of H7N9 virus with different combinations of inoculum sizes and antibody concentrations with large antibody consumption rate (dataset 2). Viral survival corresponds to antibody depletion and viral eradication coincides with antibody existence. Bifurcation diagram E) showing viral titre as a function of initial antibody concentration (schematic diagram). Viral inoculum threshold increases with increase of antibody concentration (dashed red line). Maximum capacity of viral titre remains constant if virus survives (black solid line). Purple arrows represent any viral inoculum size. A) When antibody concentration is between 0 and $A_1$, virus with any viral inoculum survive. B) and C) When antibody concentration is between $A_1$ and $A_2$ , viral kinetics survives if viral inoculum is above dashed red curve and is inhibited if viral inoculum is below dashed red curve. D) Virus with any inoculum size is inhibited when antibody concentration is greater than $A_2$. Purple, green, red, blue curve and circle represent viral kinetics with viral inoculum $10^{0.8}$, $10^{1.2}$, $10^{1.6}$ and $10^{2.0}$TCID50/ml in (A-E).

#### 3.3. Bistable kinetics of A/H1N1pdm09 with eclipse phase and small antibody consumption or large antibody consumption

In this section, we integrate H1N1pdm09 virus replication parameter (Table S2) with eclipse phase and virus neutralization parameter obtained from dataset 3 (Table S17) into System 5. Antibody-induced bistable viral kinetics exists. Due to small antibody consumption rate, antibody concentration is approximated as constant during 144 hours incubation (Fig. S23), and then initial antibody concentration is used as constant antibody concentration. Specifically, for antibody concentration $A = 0.03\, ug/ml$, virus with inoculum $10^{1.2} TCID\,50/ml$, $10^{1.6} TCID\,50/ml$ and $10^{2.0} TCID\,50/ml$ survives (Fig. S24B and Fig. S24E); for antibody concentration $A = 0.04\, ug/ml$, only virus with inoculum $10^{2.0} TCID\,50/ml$ survives (Fig. S24C and Fig. S24E). For antibody concentration less than $A_1$, virus survives independent of viral inocula (Fig. S24A and Fig. S24E). For antibody concentration higher than $A_2$, virus is inhibited independent of viral inoculum (Fig. S24D and Fig. S24E).

Next, by substituting H1N1pdm09 virus replication parameter (Table S2) with eclipse phase and virus neutralization parameter obtained from dataset 2 (Table S17) into System 5, we shown antibody-induced bistable viral kinetics also exists (Fig. S25 and Fig.S26). With initial antibody concentration $A_0 = 0.015\, ug/ml$, virus survives independent of viral inoculum and antibody is depleted (Fig. S25A and Fig. S26A). With initial antibody concentration $A_0 = 0.023\, ug/ml$, virus with viral inoculum $10^{1.2} TCID\, 50/ml$, $10^{1.6} TCID\, 50/ml$ and $10^{2.0} TCID\, 50/ml$ converges to same maximum capacity; meanwhile corresponding antibody concentration is depleted (Fig. S25B and Fig. S26B). With initial antibody concentration $A_0 = 0.027\, ug/ml$, only virus with viral inoculum $10^{2.0} TCID\, 50/ml$ survives and converges to same maximum capacity (Fig.S25C and Fig.S26C). With initial antibody concentration $A_0 = 0.03\, ug/ml$, virus is inhibited independent of viral inoculum and antibody remains (Fig. S25D and Fig.S26D).

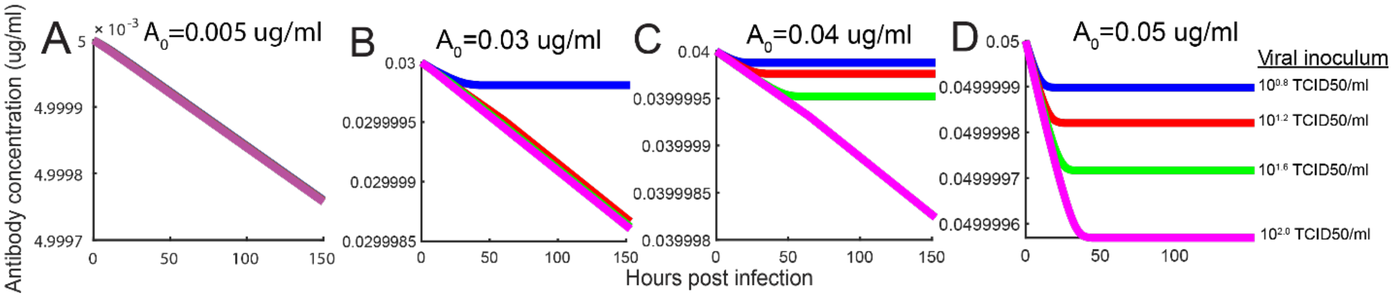


**Figure S23**. Antibody concentration kinetics with small antibody consumption rate. A-D) represents antibody concentration kinetics with initial antibody concentration $0.005\, ug/ml$, $0.03\, ug/ml$, $0.04\, ug/ml$ and $0.05\, ug/ml$.

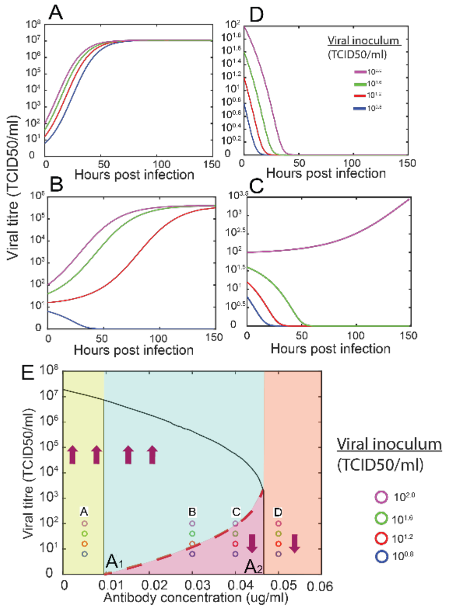


**Figure S24**. Simulated kinetics of H1N1pdm09 virus (with eclipse phase) with different combinations of inoculum sizes and antibody concentrations with small antibody consumption rate (dataset 3). Bifurcation diagram E) showing viral titre as a function of antibody concentration. Viral inoculum threshold increases with increase of antibody concentration (red dashed curve). The maximal capacity of viral titre decreases with increases of antibody concentration (black solid curve). Purple arrows represent any viral inoculum size. A) When antibody concentration is between 0 and $A_1$ virus with any viral inoculum survive. D) Virus with any inoculum size is inhibited when antibody concentration is greater than $A_2$. B) and C) When antibody concentration is between $A_1$ and $A_2$ virus survives if viral inoculum is above dashed red curve and is inhibited if viral inoculum is below dashed red curve. Purple, green, red, blue curve and circle represents viral kinetics with viral inoculum $10^{0.8}$, $10^{1.2}$, $10^{1.6}$ and $10^{2.0}$TCID50/ml in (A-E).

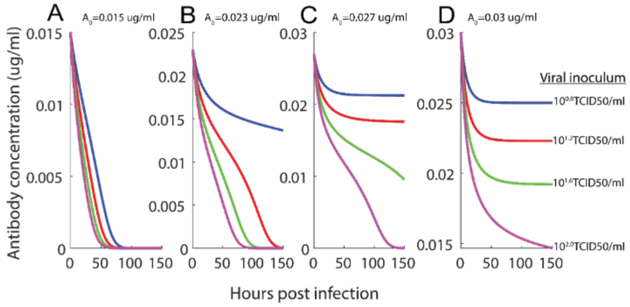


**Figure S25**. Antibody concentration kinetics (H1N1pdm09 with eclipse phase) with large antibody consumption rate. A-D) represents antibody concentration kinetics with initial antibody concentration $0.015\,ug/ml$, $0.023\,ug/ml$, $0.027$ and $0.03\,ug/ml$.

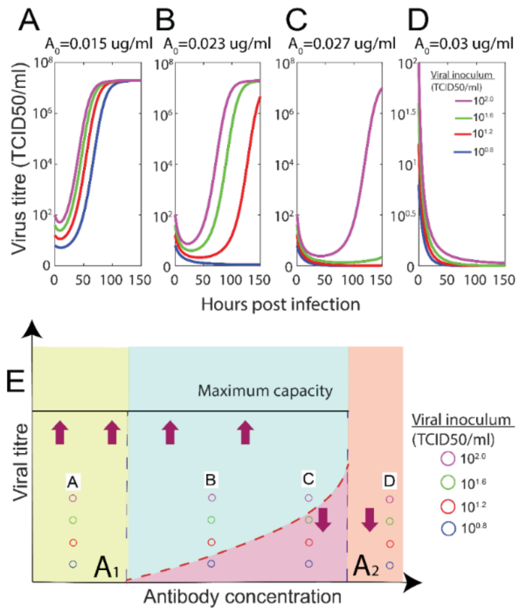


**Figure S26.** Simulated kinetics of H1N1pdm09 virus with different combinations of inoculum sizes and antibody concentrations with large antibody consumption rate (dataset 2). Viral survival corresponds to antibody depletion and viral eradication coincides with antibody existence. Bifurcation diagram E) showing viral titre as a function of initial antibody concentration (schematic diagram). Viral inoculum threshold increases with increase of antibody concentration (dashed red line). Maximum capacity of viral titre remains constant if virus survives (black solid line). Purple arrows represent any viral inoculum size. A) When antibody concentration is between 0 and $A_1$, virus with any viral inoculum survive. B) and C) When antibody concentration is between $A_1$ and $A_2$, viral kinetics survives if viral inoculum is above dashed red curve and is inhibited if viral inoculum is below dashed red curve. D) Virus with any inoculum size is inhibited when antibody concentration is greater than $A_2$. Purple, green, red, blue curve and circle represent viral kinetics with viral inoculum $10^{0.8}$, $10^{1.2}$, $10^{1.6}$ and $10^{2.0}$TCID50/ml in (A-E).

#### 3.4. Bistable kinetics of A/H7N9 with eclipse phase and small antibody consumption or large antibody consumption

In this section, we integrate H7N9 virus replication parameter (Table S2) with eclipse phase and virus neutralization parameter from dataset 3 (Table S17) into System 5. Antibody-induced bistable viral kinetics exists. Due to small antibody consumption rate, antibody concentration is approximated as constant during 144 hours incubation (Fig. S27), and then initial antibody concentration is used as constant antibody concentration. Specifically, for antibody concentration $A = 0.06\, ug/ml$, virus with inoculum $10^{1.2} TCID\, 50/ml$, $10^{1.6} TCID\, 50/ml$and $10^{2.0} TCID\, 50/ml$ survives (Fig. S28B and Fig. S28E) ; for antibody concentration $A = 0.07\, ug/ml$, only virus with inoculum $10^{2.0} TCID\, 50/ml$ survives (Fig. S28C and Fig. S28E). For antibody concentration less than $A_1$, virus survives independent of viral inoculum (Fig.S28A and Fig. S28E). For antibody concentration higher than $A_2$, virus is inhibited independent of viral inoculum (Fig.S28D and Fig. S28E).

Next, by substituting H7N9 virus replication parameter (Table S2) with eclipse phase and virus neutralization parameter obtained from dataset 2 (Table S17) into System 5, we demonstrate antibody-induced bistable viral kinetics exists (Fig. S29 and Fig. S30). With initial antibody concentration $A_0 = 0.02\, ug/ml$, virus survives independent of viral inoculum and antibody is depleted (Fig. S29A and Fig.S30A). With initial antibody concentration $A_0 = 0.041\, ug/ml$, virus with viral inoculum $10^{1.2} TCID\,50/ml$, $10^{1.6} TCID\,50/ml$ and $10^{2.0} TCID\,50/ml$ converges to same maximum capacity; meanwhile corresponding antibody concentration is depleted (Fig. S29B and S30B). With initial antibody concentration $A_0 = 0.048\, ug/ml$, only virus with viral inoculum $10^{2.0} TCID\,50/ml$ survives and converges to maximum capacity (Fig. S29C and Fig. S30C). With initial antibody concentration $A_0 = 0.055\, ug/ml$, virus is inhibited independent of viral inoculum and antibody remains (Fig. S29D and S30D).

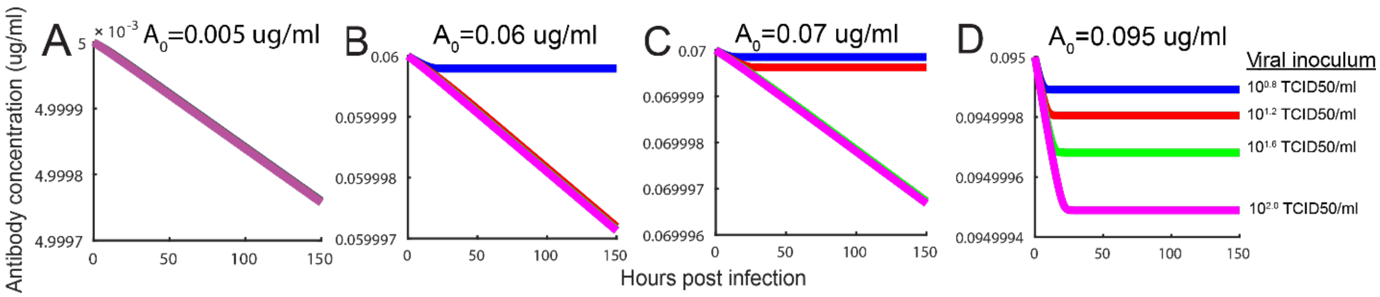


**Figure S27.** Antibody concentration kinetics (H7N9) with small antibody consumption rate. A–D) represent antibody concentration with different initial antibody concentration $0.005\, ug/ml$, $0.06\, ug/ml$, $0.07\, ug/ml$ and $0.095\, ug/ml$.

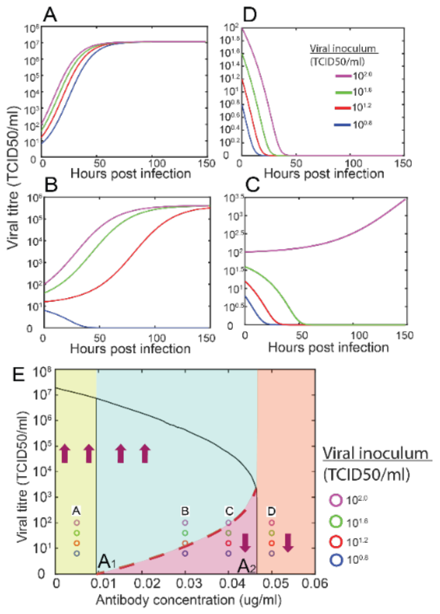


**Figure S28**. Simulated kinetics of H7N9 virus (with eclipse phase) with different combinations of inoculum sizes and antibody concentrations with low antibody consumption (dataset 3). Bifurcation diagram E) showing viral titre as a function of antibody concentration. Viral inoculum threshold increases with increase of antibody concentration (red dashed curve). The maximal capacity of viral titre decreases with increases of antibody concentration (black solid curve). Purple arrows represent any viral inoculum size. A) When antibody concentration is between 0 and $A_1$ virus with any viral inoculum survive. D) Virus with any inoculum size is inhibited when antibody concentration is greater than $A_2$. B) and C) When antibody concentration is between $A_1$ and $A_2$ virus survives if viral inoculum is above dashed red curve and is inhibited if viral inoculum is below dashed red curve. Purple, green, red, blue curve and circle represent viral kinetics with viral inoculum $10^{0.8}$, $10^{1.2}$, $10^{1.6}$ and $10^{2.0}$TCID50/ml in (A-E).

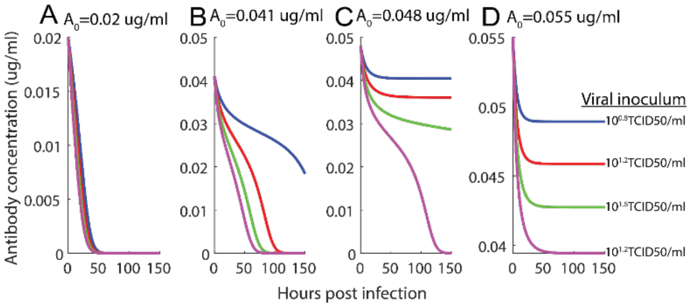


**Figure S29.** Antibody concentration kinetics with different inoculum sizes in 144-hour incubation. A–D) represents antibody concentration kinetics with initial antibody concentration $0.02\,ug/ml$, $0.041\,ug/ml$, $0.048\,ug/ml$ and $0.055\,ug/ml$.

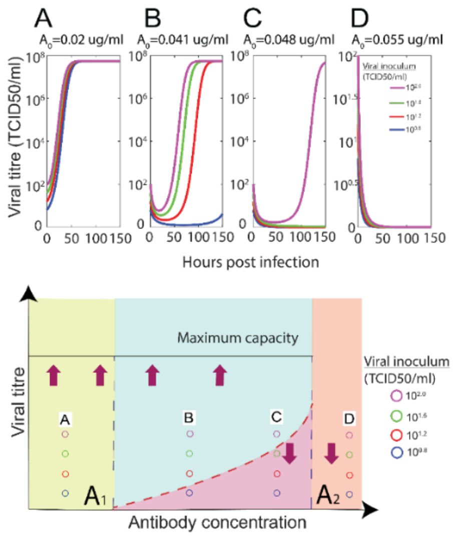


**Figure S30.** Simulated kinetics of H7N9 (virus with different combinations of inoculum sizes and antibody concentrations with large antibody consumption rate (dataset 2). Viral survival corresponds to antibody depletion and viral eradication coincides with antibody existence. Bifurcation diagram E) showing viral titre as a function of initial antibody concentration (schematic diagram). Viral inoculum threshold increases with increase of antibody concentration (dashed red line). Maximum capacity of viral titre remains constant if virus survives (black solid line). Purple arrows represent any viral inoculum size. A) When antibody concentration is between 0 and $A_1$, virus with any viral inoculum survive. B) and C) When antibody concentration is between $A_1$ and $A_2$ , viral kinetics survives if viral inoculum is above dashed red curve and is inhibited if viral inoculum is below dashed red curve. D) Virus with any inoculum size is inhibited when antibody concentration is greater than $A_2$. Purple, green, red, blue curves and circle represent viral kinetics with viral inoculum $10^{0.8}$, $10^{1.2}$, $10^{1.6}$ and $10^{2.0}$TCID50/ml in (A-D).

### 4. Viral kinetics and antibody concentration kinetics with unsaturated virus neutralization

In this section, by combining virus replication (Section.1, *Supplementary material*) and neutralization (Section.2, *Supplementary material*), we shown that for small antibody consumption, neutralizing antibody only induces monostable viral kinetics; for large antibody consumption, neutralizing antibody induce bistable viral kinetics, but its antibody concentration interval is relatively same. This result holds independent of selection of initial viral titre. Here, we illustrate that most diluted antibody first and cell control as total viral titre next.

#### 4.1. Virus neutralization parameter estimated from viral titre with the most diluted antisera as total viral titre

Below, we use virus neutralization parameter estimated from viral titre with the most diluted antisera as total viral titre. We combine A/H7N9 viral replication and virus neutralization obtained from unsaturated virus neutralization to show antibody concentration and virus kinetics. We show virus kinetics and its corresponding bifurcation diagram in Main Text and provide that antibody concentration kinetics in following section.

##### *4.1.1. Antibody concentration kinetics of A/H7N9 viral replication in presence of antibody*

First, we substitute H7N9 virus replication (Table S1) and neutralization parameter with **small antibody consumption rate** into System 4 (main text). Due to small antibody consumption rate, the amount of consumed antibody is very small and then viral kinetics with decreasing antibody concentration is approximated by viral kinetics with constant antibody concentration (Fig. S31).

Then, we substitute H7N9 virus replication (Table S1) and neutralization parameter with **large antibody consumption rate** into System 4 (main text). Due to large antibody consumption rate, viral survival corresponds to depletion of antibody; viral eradication coincides with antibody remaining, shown in Fig. S32 and Fig. 6 (main text).

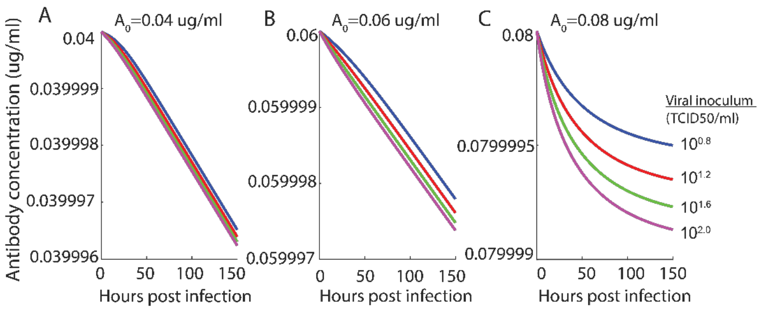


**Figure S31**. Antibody concentration kinetics with different inoculum sizes in 144-hour incubation. A-C) represents different initial antibody concentration 0.04ug/ml, 0.06ug/ml and 0.08ug/ml.

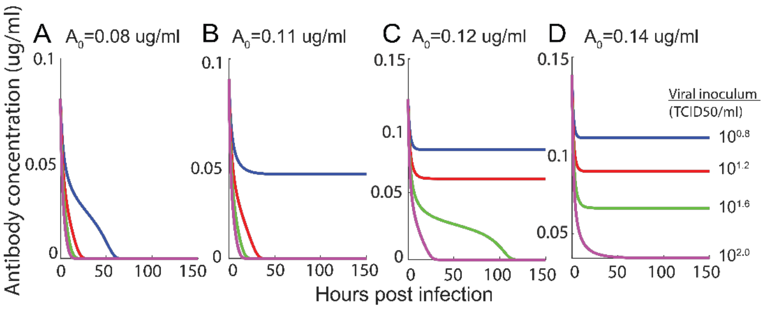


**Figure S32**. Antibody concentration kinetics with different inoculum sizes in 150-hour incubation. A-D) represents different initial antibody concentration 0.08ug/ml, 0.11ug/ml, 0.12ug/ml and 0.14ug/ml.

#### 4.2. Virus neutralization parameter estimated from viral titre with cell control as total viral titre

Below, we use virus neutralization parameter estimated from cell control with the most diluted antisera as total viral titre. We combine A/H1N1pdm09 viral replication and virus neutralization obtained from unsaturated virus neutralization to show antibody concentration and virus kinetics in following section.

##### 4.2.1. *Monostable kinetics of A/H1N1pdm09 with small antibody consumption rate*

Substituting A/H1N1pdm09 virus replication and neutralization parameter from dataset 3 with **small antibody consumption rate** (Table S17) into System 4 in main text, we shown viral kinetics only exhibit monostability and antibody concentration kinetics does not decease significantly during 144-hour incubation. Because of small antibody consumption rate, the amount of consumed antibody is very small and then viral kinetics with decreasing antibody concentration can be approximated by viral kinetics with constant antibody concentration. Antibody concentration kinetics in 144-

hour incubation is provided in Fig. S33. Then, we use initial antibody concentration as the constant antibody concentration during 144-hour incubation. Then, only one threshold $A_1$ exists to divide antibody concentration interval into two intervals (Fig. S34D). For antibody concentration less than $A_1$, virus survives independent of viral inocula (Fig. S34A, Fig. S34B and Fig. S34D). Moreover, the maximal capacity of viral capacity decrease with increase of antibody concentrations. Specifically, for antibody concentration $A = 0.005\,ug/ml$ and $A = 0.013\,ug/ml$, virus survives independent of viral inoculum size. For antibody concentration higher than $A_1$, virus is inhibited independent of viral inoculum (Fig. S34C and Fig. S34D).

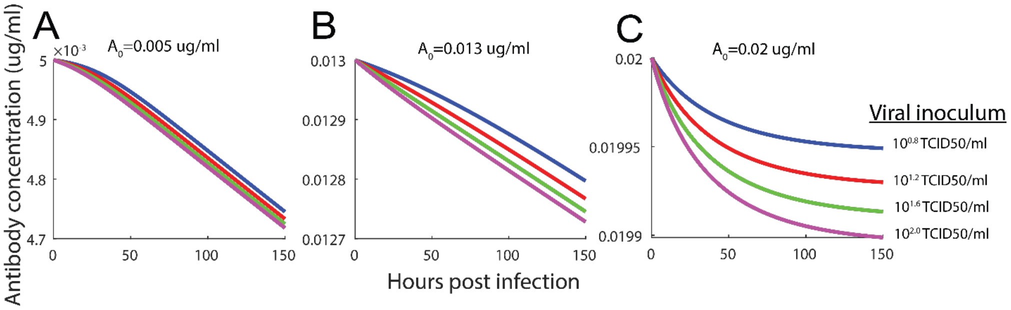


**Figure S33.** Antibody concentration kinetics with different inoculum sizes in 144-hour incubation. A–C) represent antibody concentration kinetics with different initial antibody concentration $0.005\,ug/ml$ , $0.013\,ug/ml$ and $0.02\,ug/ml$

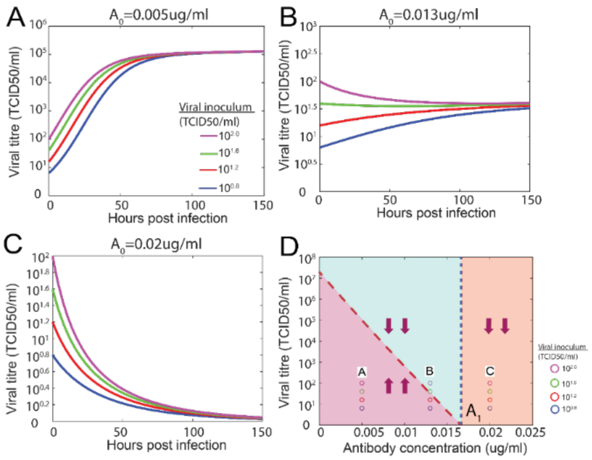


**Figure S34.** Simulated kinetics of A/H1N1pmd09 virus with different combinations of inoculum sizes and antibody concentrations with small antibody consumption (dataset 3). Bifurcation diagram D) showing viral titre as a function of initial antibody concentration. Maximal capacity of viral titre decreases with increase of antibody concentration (dashed red line). Purple arrows represent any viral inoculum size. A) and B) When antibody concentration is between 0 and $A_1$, virus with any viral inoculum survive. C) Virus with any inoculum size is inhibited when antibody concentration is greater than $A_1$. Purple, green, red, blue curve and circle represent viral kinetics with viral inoculum $10^{0.8}$, $10^{1.2}$, $10^{1.6}$ and $10^{2.0}$TCID50/ml in (A-D).

###### 4.2.2. *Bistable kinetics of A/H1N1pdm09 with large antibody consumption rate*

Substituting A/H1N1pdm09 virus replication and neutralization parameter from dataset 2 with **large antibody consumption rate** (Table S17) into System 4 in main text, we show that the viral kinetics exhibit bistability (Fig. S33 and Fig. S34), but antibody concentration interval corresponding to bistability is relatively small (Fig. S34).

With initial antibody concentration $A_0 = 0.015\, ug/ml$, virus survives independent of viral inoculum and antibody is depleted (Fig. S33A and Fig.S34A). With initial antibody concentration $A_0 = 0.03\, ug/ml$, virus with viral inoculum $10^{1.6} TCID\, 50/ml$ and $10^{2.0} TCID\, 50/ml$ survive; meanwhile corresponding antibody concentration is depleted (Fig. S33B and S34B). With initial antibody concentration $A_0 = 0.032\, ug/ml$, only virus with viral inoculum $10^{2.0} TCID\, 50/ml$ (Fig. S33C and Fig. S34C). With initial antibody concentration $A_0 = 0.035\, ug/ml$, virus is inhibited independent of viral inoculum and antibody remains (Fig. S33D and S34D).

Below, we introduce a two-dimensional deterministic model to describe viral kinetics and antibody concentration kinetics with unsaturated virus neutralization and eclipse phase, $\begin{cases} \frac{dV(t)}{dt} = \frac{\rho V(t-\tau)}{1+\beta V(t)} - \sigma V(t) - \alpha A(t)V(t) \\ \frac{dA(t)}{dt} = -\varphi A(t)V(t) \end{cases}$, (6).

By incorporating virus replication parameter (A/H1N1pdm09 and A/H7N9, Table. S1) and virus neutralization parameter (Table. S17, Table. S21) into System 4 in the main text, monostable viral kinetics exists for small antibody consumption and bistable viral kinetics exists for large antibody consumption. Moreover, by substituting virus neutralization (A/H1N1pdm09 and A/H7N9, Table. S2) and virus neutralization parameter (Table. S17, Table. S21) into System 6, monostable viral kinetics exists for small antibody consumption and bistable viral kinetics exists for large antibody consumption (data not shown).

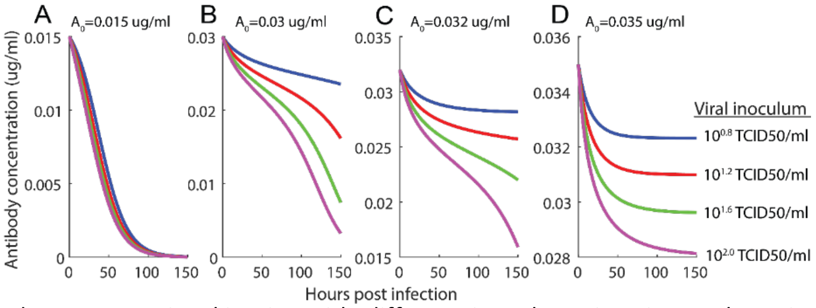


**Figure S35.** Antibody concentration kinetics with different inoculum sizes in 144-hour incubation. A–D) represent different initial antibody concentration$0.015\ ug/ml$,$0.03\ ug/ml$, $0.032\ ug/ml$ and $0.035\ ug/ml$.

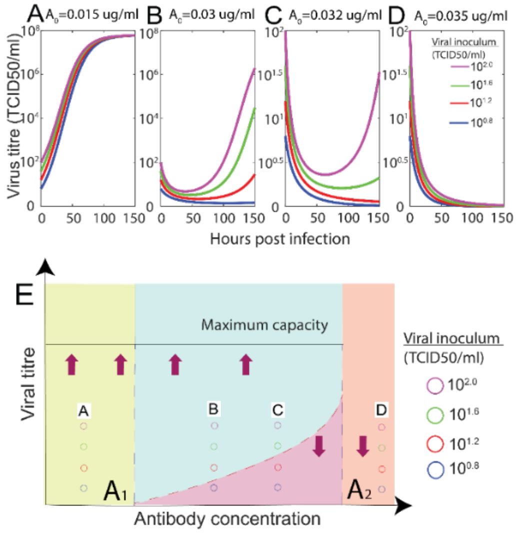


**Figure S36.** Simulated kinetics of A/H1N1pmd09 (virus with different combinations of inoculum sizes and antibody concentrations with large antibody consumption rate (dataset 2). Viral survival corresponds to antibody depletion and viral eradication coincides with antibody existence. Bifurcation diagram E) showing viral titre as a function of initial antibody concentration (schematic diagram). Viral inoculum threshold increases with increase of antibody concentration (dashed red line). Maximum capacity of viral titre remains constant if virus survives (black solid line). Purple arrows represent any viral inoculum size. A) When antibody concentration is between 0 and $A_1$, virus with any viral inoculum survive. B) and C) When antibody concentration is between $A_1$ and $A_2$ , viral kinetics survives if viral inoculum is above dashed red curve and is inhibited if viral inoculum is below dashed red curve. D) Virus with any inoculum size is inhibited when antibody concentration is greater than $A_2$. Purple, green, red, blue curves and circle represent viral kinetics with viral inoculum $10^{0.8}$, $10^{1.2}$, $10^{1.6}$ and $10^{2.0}$TCID50/ml in (A-D).

#### 5. **GISAID accession numbers and acknowledgements to generating labs**

In this section, 35 test H3N2 influenza virus with HA sequence and eight reference are shown in Table 35 and Table 36.

**Table 35.** 35 tested H3N2 influenza virus with HA sequence

| Virus name | Virus medium | GISAID_EPI | HA clade |
|---|---|---|---|
| A/Newcastle/82/2018 | MDCK-SIAT2 | EPI_ISL_395033 | 3C2.A1b/131K |
| A/Newcastle/82/2018 | Egg | EPI_ISL_339348 | 3C2.A1b/131K |
| A/Sydney/22/2018 | MDCK-SIAT,SIAT3 | EPI_ISL_363641 | 3C2.A1b/135N |
| A/Sydney/1010/2019 | MDCK-SIAT2 | EPI_ISL_363290 | 3C2.A1b/131K |
| A/Victoria/653/2017 | MDCK-SIAT1 | EPI_ISL_277940 | 3C2.A1b/135K |
| A/Victoria/653/2017 | Egg | EPI_ISL_291272 | 3C2.A1b/135K |
| A/Victoria/943/2019 | MDCK-SIAT1 | EPI_ISL_363291 | 3C2.A1b/135K |
| A/Victoria/23/2019 | MDCK-SIAT2 | EPI_ISL_363256 | 3C2.A1b/131K |
| A/Switzerland/8060/2017 | MDCK-SIAT2 | EPI_ISL_331924 | 3C2.A2/re |
| A/Switzerland/8060/2017 | Egg | EPI_ISL_331220 | 3C2.A2/re |
| A/Tasmania/511/2019 | MDCK-SIAT2 | EPI_ISL_365239 | 3C2.A1b/131K |
| A/Tasmania/512/2019 | MDCK-SIAT2 | EPI_ISL_363455 | 3C2.A1b/131K |
| A/Tasmania/519/2019 | MDCK-SIAT2 | EPI_ISL_363448 | 3C2.A1b/131K |
| A/Canberra/40/2019 | MDCK-SIAT2 | EPI_ISL_365247 | 3C2.A1b/131K |
| A/Canberra/61/2019 | MDCK-SIAT2 | EPI_ISL_356760 | 3C2.A1b/131K |
| A/Darwin/147/2019 | MDCK-SIAT2 | EPI_ISL_363278 | 3C2.A1b/131K |
| A/Darwin/148/2019 | MDCK-SIAT2 | EPI_ISL_363279 | 3C2.A1b/131K |
| A/Darwin/157/2019 | MDCK-SIAT2 | EPI_ISL_363281 | 3C2.A1b/131K |
| A/Darwin/158/2019 | MDCK-SIAT2 | EPI_ISL_363282 | 3C2.A1b/131K |
| A/Christchurch/514/2019 | MDCK-SIAT1 | EPI_ISL_363269 | 3C2.A1b/131K |
| A/Christchurch/516/2019 | MDCK-SIAT1 | EPI_ISL_363270 | 3C2.A1b/131K |
| A/Christchurch/518/2019 | MDCK-SIAT1 | EPI_ISL_363271 | 3C2.A1b/131K |
| A/Brunei/16/2019 | MDCK-SIAT1 | EPI_ISL_363434 | 3C2.B |
| A/Canberra/107/2019 | MDCK-SIAT1 | EPI_ISL_363205 | 3C2.A1b/131K |
| A/Canberra/108/2019 | MDCK-SIAT1 | EPI_ISL_363206 | 3C2.A1b/131K |
| A/Canberra/109/2019 | MDCK-SIAT1 | EPI_ISL_363207 | 3C2.A1b/131K |
| A/Canberra/110/2019 | MDCK-SIAT1 | EPI_ISL_363208 | 3C2.A1b/131K |
| A/Fiji/15/2019 | MDCK-SIAT1 | EPI_ISL_363437 | 3C2.A1b/131K |
| A/Fiji/25/2019 | MDCK-SIAT1 | EPI_ISL_363438 | 3C2.A1b/131K |
| A/Fiji/30/2019 | MDCK-SIAT1 | EPI_ISL_363439 | 3C2.A1b/131K |
| A/Fiji/31/2019 | MDCK-SIAT1 | EPI_ISL_363440 | 3C2.A1b/131K |
| A/Fiji/36/2019 | MDCK-SIAT1 | EPI_ISL_363441 | 3C2.A1b/131K |
| A/Hong Kong/4801/2014 | MDCK-SIAT4 | EPI_ISL_165553 | 3C2.A |
| A/Brisbane/32/2017 | Egg | EPI_ISL_363256 | 3C2.A1a |

**Table 36.** Eight reference H3N2 influenza virus

| Virus name | Virus medium | GISAID_EPI | HA clade |
|---|---|---|---|
| A/Newcastle/82/2018 | MDCK-SIAT2 | EPI_ISL_395033 | 3C2.A1b/131K |
| A/Newcastle/82/2018 | Egg | EPI_ISL_339348 | 3C2.A1b/131K |
| A/Sydney/22/2018 | MDCK-SIAT,SIAT3 | EPI_ISL_363641 | 3C2.A1b/135N |
| A/Victoria/653/2017 | MDCK-SIAT1 | EPI_ISL_277940 | 3C2.A1b/135K |
| A/Victoria/653/2017 | Egg | EPI_ISL_291272 | 3C2.A1b/135K |
| A/Switzerland/8060/2017 | MDCK-SIAT2 | EPI_ISL_331924 | 3C2.A2/re |
| A/Switzerland/8060/2017 | Egg | EPI_ISL_331220 | 3C2.A2/re |
| A/Hong Kong/4801/2014 | MDCK-SIAT4 | EPI_ISL_165553 | 3C2.A |

Supplementary Material

*We acknowledge the authors, originating and submitting laboratories of the sequences from GISAID's EpiFlu™ Database on which this research is based. The list is detailed below.*

*All submitters of data may be contacted directly via the GISAID website* ***www.gisaid.org***

| Segment ID | Segment | Country | Collection date | Isolate-ID | Isolate name | Originating Lab | Submitting Lab | Authors |
|---|---|---|---|---|---|---|---|---|
| EPI1484756 | HA | Brunei | 2019-Mar-25 | EPI_ISL_363434 | A/Brunei/16/2019 | R.I.P.A.S. Hospital, Department of Laboratory Services | WHO Collaborating Centre for Reference and Research on Influenza | Deng,Y-M; Iannello,P; Lau,H; Todd,A; Spirason,N; Komadina,N. |
| EPI1060375 | HA | Australia | 2017-Aug-10 | EPI_ISL_277940 | A/Victoria/653/2017 | Monash Medical Centre | WHO Collaborating Centre for Reference and Research on Influenza | Deng,Y-M; Iannello,P; Lau,H; Kaye,M; Todd,A; Komadina,N |
| EPI1140329 | HA | Australia | 2017-Aug-10 | EPI_ISL_291272 | A/Victoria/653/2017 | Monash Medical Centre | WHO Collaborating Centre for Reference and Research on Influenza | Deng,Y-M.; Iannello,P.; Lau,H.; Kaye,M.; Todd,A.; Komadina,N. |
| EPI1639383 | HA | Australia | 2019-Apr-27 | EPI_ISL_363208 | A/Canberra/110/2019 | Canberra Hospital | WHO Collaborating Centre for Reference and Research on Influenza | Deng,Y-M; Iannello,P; Lau,H; Todd,A; Spirason,N; Komadina,N. |
| EPI1484388 | HA | Australia | 2019-Apr-24 | EPI_ISL_363205 | A/Canberra/107/2019 | Canberra Hospital | WHO Collaborating Centre for Reference and Research on Influenza | Deng,Y-M; Iannello,P; Lau,H; Todd,A; Spirason,N; Komadina,N. |
| EPI1639462 | HA | Australia | 2019-Apr-04 | EPI_ISL_363291 | A/Victoria/943/2019 | Royal Chidrens Hospital | WHO Collaborating Centre for Reference and Research on Influenza | Deng,Y-M; Iannello,P; Lau,H; Todd,A; Spirason,N; Komadina,N. |
| EPI1639450 | HA | Australia | 2019-Apr-10 | EPI_ISL_363279 | A/Darwin/148/2019 | Influenza Surveillance Centre for Disease Control | WHO Collaborating Centre for Reference and Research on Influenza | Deng,Y-M; Iannello,P; Lau,H; Todd,A; Spirason,N; Komadina,N. |
| EPI1484390 | HA | Australia | 2019-Apr-26 | EPI_ISL_363207 | A/Canberra/109/2019 | Canberra Hospital | WHO Collaborating Centre for Reference and Research on Influenza | Deng,Y-M; Iannello,P; Lau,H; Todd,A; Spirason,N; Komadina,N. |
| EPI1639452 | HA | Australia | 2019-Apr-14 | EPI_ISL_363281 | A/Darwin/157/2019 | Influenza Surveillance Centre for Disease Control | WHO Collaborating Centre for Reference and Research on Influenza | Deng,Y-M; Iannello,P; Lau,H; Todd,A; Spirason,N; Komadina,N. |
| EPI1639449 | HA | Australia | 2019-Apr-08 | EPI_ISL_363278 | A/Darwin/147/2019 | Influenza Surveillance Centre for Disease Control | WHO Collaborating Centre for Reference and Research on Influenza | Deng,Y-M; Iannello,P; Lau,H; Todd,A; Spirason,N; Komadina,N. |
| EPI1639382 | HA | Australia | 2019-Apr-26 | EPI_ISL_363206 | A/Canberra/108/2019 | Canberra Hospital | WHO Collaborating Centre for Reference and Research on Influenza | Deng,Y-M; Iannello,P; Lau,H; Todd,A; Spirason,N; Komadina,N. |
| EPI1639453 | HA | Australia | 2019-Apr-14 | EPI_ISL_363282 | A/Darwin/158/2019 | Influenza Surveillance Centre for Disease Control | WHO Collaborating Centre for Reference and Research on Influenza | Deng,Y-M; Iannello,P; Lau,H; Todd,A; Spirason,N; Komadina,N. |
| EPI1639442 | HA | New Zealand | 2019-Apr-08 | EPI_ISL_363271 | A/Christchurch/518/2019 | Canterbury Health Services | WHO Collaborating Centre for Reference and Research on Influenza | Deng,Y-M; Iannello,P; Lau,H; Todd,A; Spirason,N; Komadina,N. |
| EPI1484453 | HA | New Zealand | 2019-Apr-03 | EPI_ISL_363270 | A/Christchurch/516/2019 | Canterbury Health Services | WHO Collaborating Centre for Reference and Research on Influenza | Deng,Y-M; Iannello,P; Lau,H; Todd,A; Spirason,N; Komadina,N. |
| EPI1484924 | HA | Australia | 2019-Mar-07 | EPI_ISL_363455 | A/Tasmania/512/2019 | Hobart Pathology | WHO Collaborating Centre for Reference and Research on Influenza | Deng,Y-M; Iannello,P; Lau,H; Todd,A; Spirason,N; Komadina,N. |
| EPI1639441 | HA | New Zealand | 2019-Apr-05 | EPI_ISL_363269 | A/Christchurch/514/2019 | Canterbury Health Services | WHO Collaborating Centre for Reference and Research on Influenza | Deng,Y-M; Iannello,P; Lau,H; Todd,A; Spirason,N; Komadina,N. |
| EPI1639428 | HA | Australia | 2019-Apr-26 | EPI_ISL_363256 | A/Victoria/23/2019 | Victorian Infectious Diseases Reference Laboratory | WHO Collaborating Centre for Reference and Research on Influenza | Deng,Y-M; Iannello,P; Lau,H; Todd,A; Spirason,N; Komadina,N. |
| EPI1484868 | HA | Australia | 2019-Mar-22 | EPI_ISL_363448 | A/Tasmania/519/2019 | Hobart Pathology | WHO Collaborating Centre for Reference and Research on Influenza | Deng,Y-M; Iannello,P; Lau,H; Todd,A; Spirason,N; Komadina,N. |
| EPI1492987 | HA | Australia | 2019-Mar-07 | EPI_ISL_365239 | A/Tasmania/511/2019 | Hobart Pathology | WHO Collaborating Centre for Reference and Research on Influenza | Deng,Y-M; Iannello,P; Lau,H; Todd,A; Spirason,N; Komadina,N. |
| EPI1444768 | HA | Australia | 2019-Mar-17 | EPI_ISL_356760 | A/Canberra/61/2019 | Canberra Hospital | WHO Collaborating Centre for Reference and Research on Influenza | Deng,Y-M; Iannello,P; Lau,H; Todd,A; Spirason,N; Moselen,J; Komadina,N. |
| EPI1484812 | HA | Fiji | 2019-Mar-11 | EPI_ISL_363441 | A/Fiji/36/2019 | National Centre for Scientific Services for Virology and V | WHO Collaborating Centre for Reference and Research on Influenza | Deng,Y-M; Iannello,P; Lau,H; Todd,A; Spirason,N; Komadina,N. |
| EPI1484780 | HA | Fiji | 2019-Feb-19 | EPI_ISL_363437 | A/Fiji/15/2019 | National Centre for Scientific Services for Virology and V | WHO Collaborating Centre for Reference and Research on Influenza | |
| EPI1484796 | HA | Fiji | 2019-Mar-09 | EPI_ISL_363439 | A/Fiji/30/2019 | National Centre for Scientific Services for Virology and V | WHO Collaborating Centre for Reference and Research on Influenza | Deng,Y-M; Iannello,P; Lau,H; Todd,A; Spirason,N; Komadina,N. |
| EPI1484788 | HA | Fiji | 2019-Feb-26 | EPI_ISL_363438 | A/Fiji/25/2019 | National Centre for Scientific Services for Virology and V | WHO Collaborating Centre for Reference and Research on Influenza | Deng,Y-M; Iannello,P; Lau,H; Todd,A; Spirason,N; Komadina,N. |
| EPI1639461 | HA | Australia | 2019-Apr-08 | EPI_ISL_363290 | A/Sydney/1010/2019 | Institute of Medical and Veterinary Science (IMVS) | WHO Collaborating Centre for Reference and Research on Influenza | Deng,Y-M; Iannello,P; Lau,H; Todd,A; Spirason,N; Komadina,N. |
| EPI1484804 | HA | Fiji | 2019-Mar-10 | EPI_ISL_363440 | A/Fiji/31/2019 | National Centre for Scientific Services for Virology and V | WHO Collaborating Centre for Reference and Research on Influenza | Deng,Y-M; Iannello,P; Lau,H; Todd,A; Spirason,N; Komadina,N. |
| EPI1359999 | HA | Australia | 2018-Dec-23 | EPI_ISL_339348 | A/Newcastle/82/2018 | John Hunter Hospital | WHO Collaborating Centre for Reference and Research on Influenza | Deng,Y-M; Iannello,P; Lau,H; Kaye,M; Todd,A; Spirason,N; Komadina,N. |
| EPI1493051 | HA | Australia | 2019-Feb-16 | EPI_ISL_365247 | A/Canberra/40/2019 | Canberra Hospital | WHO Collaborating Centre for Reference and Research on Influenza | Deng,Y-M; Iannello,P; Lau,H; Todd,A; Spirason,N; Komadina,N. |
| EPI1607125 | HA | Australia | 2018-Dec-23 | EPI_ISL_395033 | A/Newcastle/82/2018 | WHO Collaborating Centre for Reference and Research o | Centers for Disease Control and Prevention | |
| EPI1323470 | HA | Switzerland | 2018-Jan-01 | EPI_ISL_331924 | A/Switzerland/8060/2017 | National Institute for Medical Research | Centers for Disease Control and Prevention | |
| EPI1318863 | HA | Switzerland | 2018-Nov-01 | EPI_ISL_331220 | A/Switzerland/8060/2017 | | National Institute for Biological Standards and Control (NIBSC) | Nicolson, Carolyn |
| EPI1044843 | HA | Australia | 2017-Mar-09 | EPI_ISL_274984 | A/Brisbane/32/2017 | Queensland Health Forensic and Scientific Services | WHO Collaborating Centre for Reference and Research on Influenza | Deng,Y-M; Iannello,P; Lau,H; Kaye,M; Todd,A; Komadina,N. |
| EPI1485365 | HA | Australia | 2018-Mar-03 | EPI_ISL_363641 | A/Sydney/22/2018 | Clinical Virology Unit, CDIM | WHO Collaborating Centre for Reference and Research on Influenza | Deng,Y-M; Iannello,P; Lau,H; Todd,A; Spirason,N; Komadina,N. |
| EPI539574 | HA | Hong Kong (SAR) | 2014-Feb-26 | EPI_ISL_165553 | A/Hong Kong/4800/2014 | Government Virus Unit | National Institute for Medical Research | |
